# Postsynaptic TAK1-GluN2B axis supports hippocampal synaptic integrity and cognitive function during aging

**DOI:** 10.64898/2026.10.25.761716

**Authors:** Yang Liu, Mao-Qing Huang, Yi-Ci Zhang, Zhuo-Ran Jia, Xia Min, Zi-Yue Zhou, Jia-Yao Chen, Fang-Jiao Zong, Jing Gao, Yiwen Xu, Tong Wang, Yaoyang Zhang, Kai-Wen He

**Author notes:** Correspondence: Kai-Wen He. These authors contributed equally (Yang Liu, Mao-Qing Huang).

## Abstract

Transforming growth factor β-activated kinase 1 (TAK1) is a serine/threonine kinase best known for its roles in immune and stress-response signaling, yet its physiological function in mature neurons remains poorly understood. Here we identify neuronal TAK1 as a regulator of excitatory synaptic integrity, hippocampal plasticity and memory. Conditional reduction of TAK1 in forebrain excitatory neurons impaired short- and long-term memory and was accompanied by attenuated long-term potentiation in the hippocampus. At the cellular level, TAK1 deficiency reduced dendritic spine density and suppressed excitatory synaptic transmission, whereas genetic or pharmacological disruption of TAK1 kinase activity recapitulated these synaptic deficits

Mechanistically, TAK1 localized to the postsynaptic compartment of excitatory synapses and associated with PSD-95. Phosphoproteomic analysis of isolated postsynaptic density fractions revealed coordinated changes in synaptic phosphorylation following TAK1 reduction, including decreased phosphorylation of the NMDA receptor subunit GluN2B at serine 920. Analyses of phosphodeficient and phosphomimetic GluN2B mutants supported a role for S920 phosphorylation downstream of TAK1 in maintaining dendritic spine density, excitatory synaptic transmission and spontaneous synaptic calcium activity.

Notably, synaptic TAK1 level was reduced in the aged hippocampus. Re-expressing neuronal TAK1 in aged mice increased dendritic spine density, enhanced hippocampal synaptic plasticity, and improved memory performance. Together, these findings define a postsynaptic TAK1–GluN2B signaling axis that supports excitatory synaptic function and suggest that reduced synaptic TAK1 signaling may contribute to hippocampal vulnerability during aging.

**Graphical abstract:** 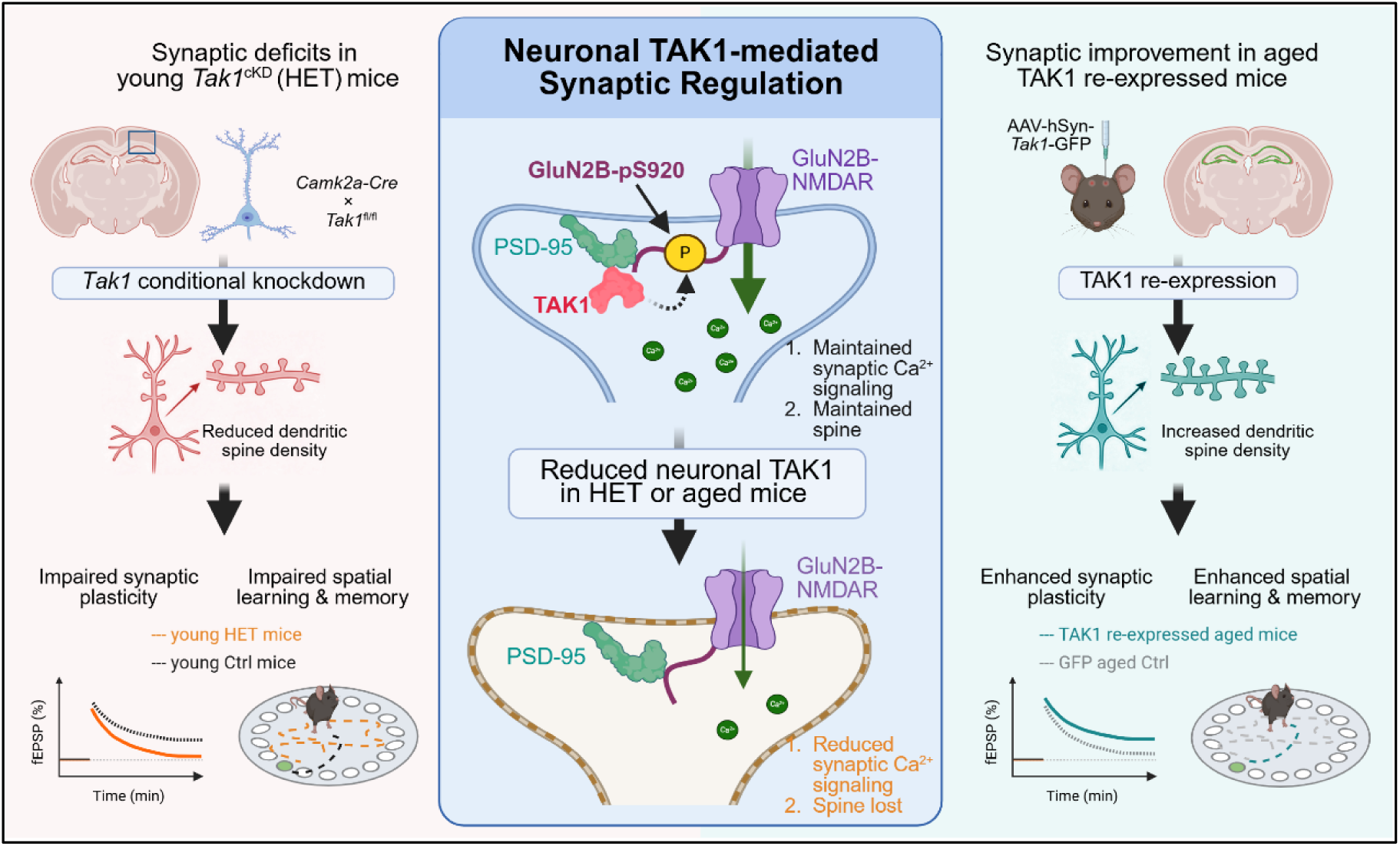

## Introduction

The preservation of excitatory synapses throughout adult life is essential for cognitive resilience. During brain aging, cognitive decline is associated less with widespread neuronal loss than with selective alterations in synaptic structure, dendritic spine morphology and synaptic efficacy in vulnerable circuits, including the hippocampus and prefrontal cortex^1–3^. These observations have led to the concept that age-related cognitive vulnerability reflects, at least in part, failure of molecular programs that maintain synaptic integrity over time. However, the signaling mechanisms that actively preserve mature excitatory synapses during physiological aging remain incompletely understood.

Excitatory synapse stability depends on highly organized postsynaptic signaling complexes assembled around glutamate receptors, scaffold proteins and actin-regulatory machinery within the postsynaptic density. Protein phosphorylation provides a rapid and reversible mechanism for coordinating these complexes by regulating receptor trafficking, cytoskeletal remodeling and calcium-dependent signaling^4–6^. Many serine/threonine kinases, including CaMKII, PKA, ERK, regulate complementary aspects of synaptic efficacy, long-term synaptic plasticity and activity-dependent structural remodeling^7–9^. NMDA receptors are particularly important in this context because they couple synaptic activity to calcium signaling programs that shape dendritic spine stability and circuit refinement. The GluN2B cytoplasmic tail is extensively regulated by phosphorylation and protein interactions that influence receptor localization, channel properties, signaling output and synaptic plasticity^10–12^. Although kinase-dependent mechanisms of acute synaptic plasticity have been intensively studied, much less is known about phosphorylation pathways that sustain excitatory synapse integrity across adult life and aging^1,2,13^.

Transforming growth factor-β-activated kinase 1 (TAK1; *Map3k7 or Tak1*) is a MAP kinase kinase kinase best known as a signaling hub downstream of inflammatory cytokines, innate immune receptors and cellular stress pathways^14,15^. Consequently, studies of TAK1 in the nervous system have focused primarily on neuroinflammation, neuronal injury and cell-death signaling^16–18^. Nevertheless, large-scale transcriptomic datasets indicate that Map3k7 is broadly expressed across adult brain cell types, including excitatory neurons^19,20^, raising the possibility that TAK1 may also have physiological neuronal functions. Consistent with this possibility, TAK1 was identified in Drosophila as an essential regulator of presynaptic homeostatic plasticity^21^. In addition, TAK1 protein levels decline in aged human brain^18^, suggesting a potential relationship between TAK1 loss and age-associated neuronal vulnerability. These observations raise an important but unresolved question: whether TAK1 functions at mammalian central synapses, and whether age-associated reduction of TAK1 contributes directly to deterioration of postsynaptic integrity.

Here we identify neuronal TAK1 as a postsynaptic regulator of excitatory synapse maintenance. We show that selective reduction of TAK1 in forebrain excitatory neurons disrupts dendritic spine density, alters excitatory synaptic transmission, impairs hippocampal long-term potentiation and compromises hippocampus-dependent memory. Mechanistically, TAK1 localizes to the postsynaptic compartment, associates with PSD-95-containing signaling complexes and maintains phosphorylation of the NMDA receptor subunit GluN2B at serine 920, thereby supporting spontaneous synaptic calcium signaling and spine integrity. Finally, we show that synaptic TAK1 is reduced in the aged hippocampus and that re-expressing neuronal TAK1 in aged mice improves dendritic spine density, hippocampal plasticity and memory performance. These findings define a postsynaptic TAK1–GluN2B signaling pathway that may support the excitatory synapse integrity and cognitive function during aging.

## Results

### Neuronal TAK1 is required for hippocampal synaptic plasticity and memory

To determine the physiological function of neuronal TAK1 in the young adult brain, we generated forebrain excitatory neuron-specific TAK1 conditional heterozygous knockout mice by crossing *Camk2a-Cre* mice with *Tak1* floxed mice, thereby selectively reducing TAK1 expression in postnatal excitatory neurons (Fig. 1A). Heterozygous conditional knockout (*Camk2a-Cre*; *Tak1*^fl/+^, HET) mice were viable, exhibited normal body weight (Fig. 1B) and showed no detectable changes in the hippocampal cytokines examined (Fig. S1A, B). Hippocampal *Tak1* mRNA was modestly reduced (Fig. 1C), consistent with partial depletion of neuronal TAK1.

**Figure 1.**
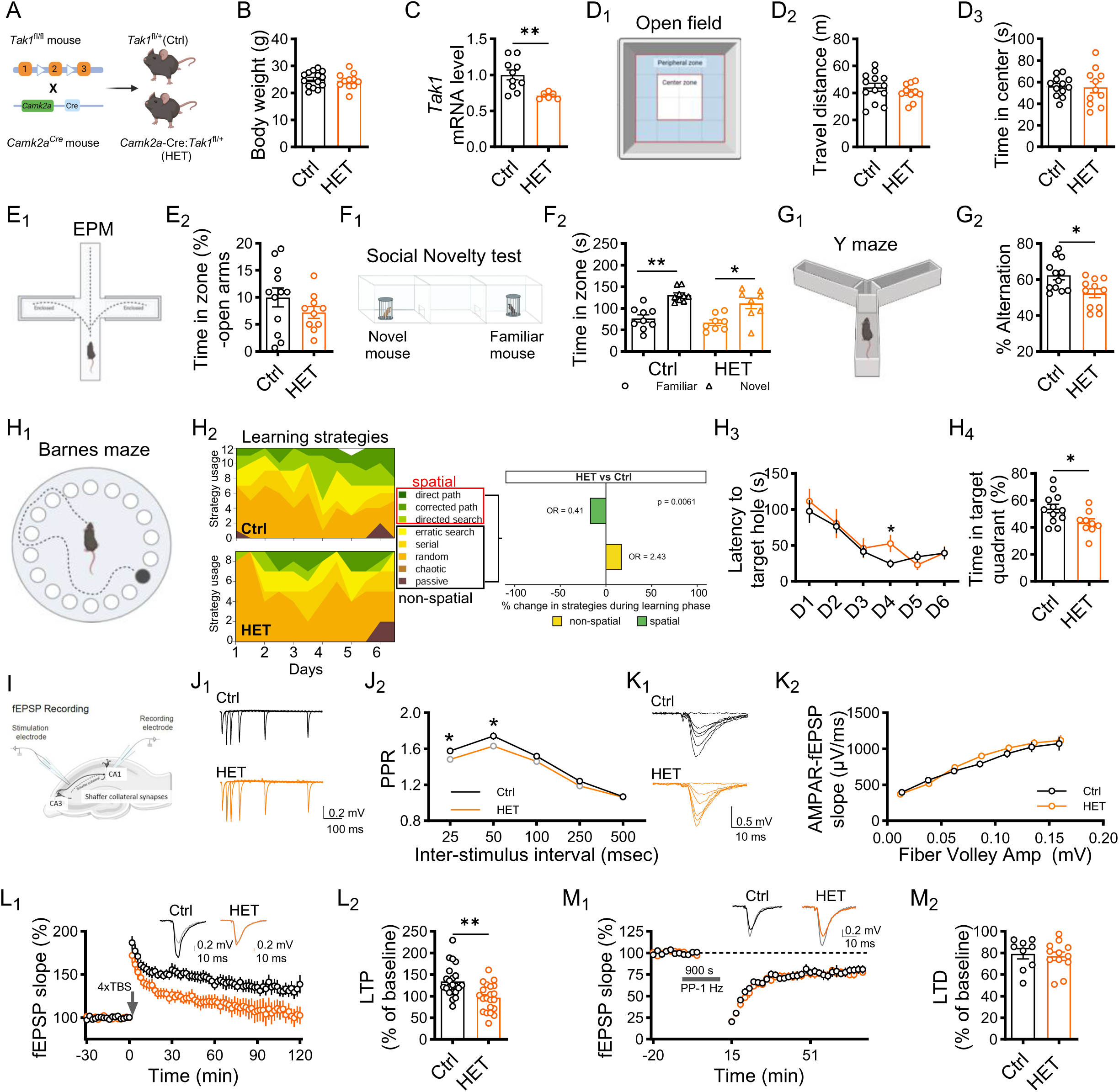
Neuronal TAK1 is required for hippocampal synaptic plasticity and memory. **A**, Generation of forebrain excitatory neuron-specific Tak1 conditional knockdown (*Camk2a*-cre; *Tak1*^fl/+^, HET) mice by crossing Camk2a-Cre mice with Tak1 floxed mice. **B**, Body weight of adult control (*Tak1*^fl/+^, Ctrl) and HET mice (Ctrl: n = 14; HET: n = 11P = 0.768). **C**, Comparison of hippocampal *Tak1* mRNA expression between HET and Ctrl mice (Ctrl: n = 10; HET: n = 6; P = 0.003). **D**, Open-field test diagram (**D_1_**) and comparison of total travel distance (**D_2_**) and time spent in the center zone (**D_3_**) between HET and Ctrl. (Ctrl: n = 13; HET: n = 10). **E**, Elevated plus maze diagram (**E_1_**) and comparison of the percentage of time spent in the open arms between groups (**E_2_**). (Ctrl: n = 12; HET: n = 10). **F**, Three-chamber social-interaction test assay diagram (**F_1_**) and comparison of the time spent investigating the novel versus familiar mouse between groups (**F_2_**). (Ctrl: n = 9, P = 0.004; HET: n = 8, P = 0.016; Wilcoxon matched-pairs signed rank test). **G**, Y-maze diagram (**G_1_**) and comparison of the spontaneous alternation performance between groups (**G_2_**). (Ctrl: n = 12; HET: n = 11; P = 0.046). **H**, Barnes maze (Ctrl: n = 12; HET: n = 9). (**H_1_**), experimental paradigm. (**H_2_**), search strategy analysis during acquisition training (P = 0.006). (**H_3_**), escape latency across training days (P_D4 = 0.032, unpaired t test). (**H_4_**), percentage of time spent in the target quadrant during the probe trial (P = 0.041). **I**, Diagram demonstrating the fEPSP recording from Schaffer collateral–CA1 synapses. **J**, Comparison of paired-pulse ratio of the Schaffer collateral–CA1 synapses between HET and Ctrl mice (Ctrl: n = 31 slices from 4 mice; HET: n = 31 slices from 4 mice). (**J_1_**), Representative traces. (**J_2_**), Paired-pulse ratio measured at indicated interstimulus intervals (P_25ms_ = 0.019; P_50ms_ = 0.044; unpaired t-test with Welch correction). **K**, Comparison of basal excitatory synaptic transmission (Ctrl: n = 36 slices from 5 mice; HET: n = 32 slices from 4 mice). (**K_1_**), Representative fEPSP waveforms. (**K_2_**), Comparison of input-output relationship between fiber volley amplitude and fEPSP slope (unpaired t-test with Welch correction). **L**, Comparison of the TBS-induced LTP in the Schaffer collateral–CA1 synapses (Ctrl: n = 22 slices from 7 mice; HET: n = 21 slices from 6 mice). (**L_1_**), Entire time course of LTP. Top insert, representative waveforms (Gray: baseline; Black and Orange: 2 h post induction). (**L_2_**), Quantification of LTP measured during the last 10 min of recording (P = 0.001). **M**, Comparison of the paired-pulse 1 Hz-induced LTD in the Schaffer collateral–CA1 synapses (Ctrl: n = 9 slices from 4 mice; HET: n = 12 slices from 4 mice). (**M_1_**), Entire time course of LTD. Top insert, representative waveforms (Gray: baseline; Black and Orange: 1h post-induction). (**M_2_**), Quantification of LTD measured during the last 10 min of recording. Data are presented as mean ± s.e.m. Statistical tests: Two-sided Mann–Whitney U test was used unless otherwise stated. Exact P values are reported for comparisons indicated in the figure; complete statistical results, including nonsignificant comparisons, are provided in the Key resource table 4. Data summary.

We first asked whether partial neuronal TAK1 reduction affected general behavioral performance that could confound the interpretation of cognitive tests. Young adult (2-3-month-old) HET mice displayed normal locomotor activity and anxiety-related behavior in both the open-field and elevated plus maze tests (Fig. 1D, E; Fig. S1C). In the three-chamber assay, control and HET mice showed comparable preferences for a novel mouse over either an empty chamber (Fig. S1D) or a familiar mouse (Fig. 1F). These findings indicate that partial neuronal TAK1 deficiency does not produce detectable impairment in locomotion, anxiety-related behavior or social interaction.

Having established normal baseline behaviors, we next examined hippocampus-dependent learning and memory. In the Y-maze spontaneous alternation task, HET mice showed reduced spontaneous alternation (Fig. 1G) despite normal overall locomotor activity (Fig. S1E), indicating impaired spatial working memory. HET mice also exhibited deficits in the Barnes maze. During training, HET mice adopted spatial search strategies less frequently than controls (Fig. 1H_2_; Fig. S1F; see Methods) and showed slower acquisition of the task (Fig. 1H_3_). During the probe trial, HET mice spent less time in the target quadrant (Fig. 1H_4_). Together, these findings demonstrate that partial neuronal TAK1 depletion compromises hippocampus-dependent spatial learning and memory.

To determine whether these cognitive deficits were accompanied by altered hippocampal synaptic function, we performed extracellular field potential recordings at Schaffer collateral–CA1 synapses (Fig. 1I). HET mice exhibited a reduced paired-pulse ratio at interstimulus intervals of 25–50 ms (Fig. 1J), consistent with an increase in presynaptic release probability and with the previously reported role of TAK1 in presynaptic regulation^21^. Despite this alteration, input–output relationships were comparable between genotypes (Fig. 1K), indicating that fast basal synaptic transmission was largely preserved.

Long-term synaptic plasticity provides a principal cellular substrate for learning and memory. We therefore examined whether neuronal TAK1 deficiency differentially affects long-term potentiation (LTP) and long-term depression (LTD). Theta-burst stimulation-induced LTP was impaired in HET mice relative to littermate controls (Fig. 1L), whereas paired-pulse 1 Hz stimulation-induced LTD remained largely intact (Fig. 1M). The selective impairment of LTP closely parallels the deficits in hippocampus-dependent spatial memory observed in HET mice.

Together, these findings demonstrate that neuronal TAK1 is required for normal hippocampal synaptic plasticity and hippocampus-dependent memory.

### Neuronal TAK1 supports excitatory synapse structure and function

The impaired LTP and spatial memory observed in HET mice raised the possibility that neuronal TAK1 may contribute to the maintenance of excitatory synapses. To directly examine the cell-autonomous role of TAK1, we acutely reduced TAK1 expression in cultured hippocampal neurons using shRNA-mediated knockdown. Primary neurons were transfected at DIV8 and analyzed at DIV16 by morphological imaging and whole-cell electrophysiology (Fig. 2A; Fig. S2A). Among the three independent shRNAs tested, shRNA-2 most efficiently reduced both mRNA and protein levels of TAK1 without altering the cytokines examined (Fig. S2B, C), shRNA-2 was therefore used for subsequent experiments (Fig. 2B).

**Figure 2.**
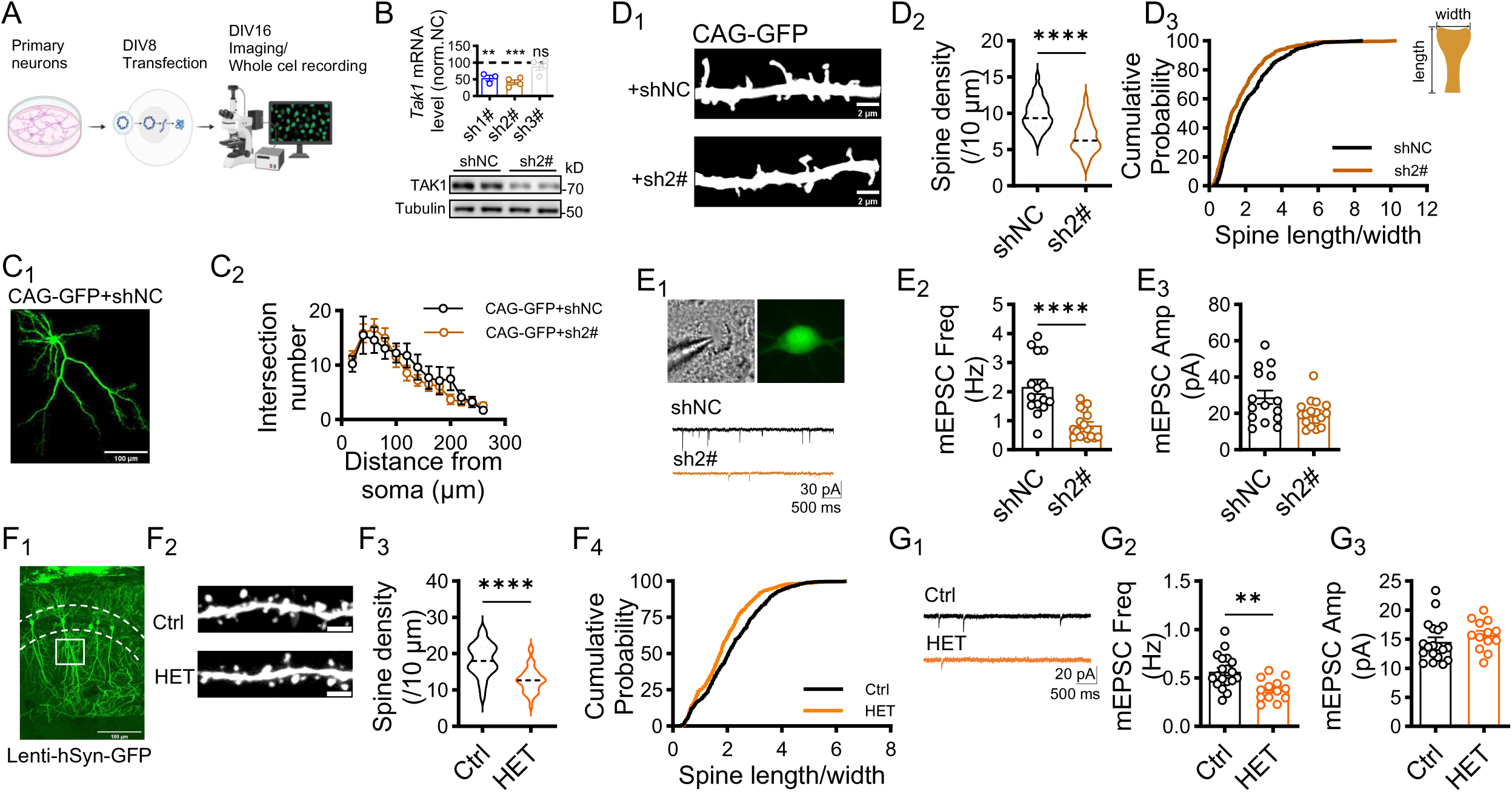
Neuronal TAK1 deficiency reduces dendritic spine density and excitatory synaptic transmission. **A**, Experimental timeline for TAK1 knockdown, morphological analysis and whole-cell recording in primary cultured neurons. **B,** Relative *Tak1* mRNA expression (upper panel) in neurons transfected with a non-targeting control shRNA (shNC) or one of three *Tak1*-targeting shRNAs (sh1#, sh2# and sh3#; n = 3, 4 or 4 wells; P_shNC vs. sh1# = 0.03; P_shNC vs. sh2# < 0.001; P_shNC vs. sh3# = 0.377). Immunoblot analysis of TAK1 and tubulin (lower panel) in primary hippocampal culture. Molecular masses are indicated in kilodaltons (kDa) on the right side. **C,** Dendritic arborization following TAK1 knockdown. (**C_1_**), Representative image of a neuron co-expressing CAG-gag-GFP (CAG-GFP) and shNC. Scale bar, 100 μm. (**C_2_**), Sholl analysis of neurons expressing CAG-GFP together with shNC or sh2# (shNC: n = 10 neurons from 2 cultures; sh2#: n = 10 neurons from 2 cultures). **D,** Dendritic spine density and morphology analysis following TAK1 knockdown. (**D_1_**), Representative dendritic segments from neurons expressing shNC or sh2#. Scale bars, 2 μm. (**D_2_**), Quantification of dendritic spine density (shNC: n = 33 dendritic segments from 17 neurons from 2 cultures; sh2#: n = 28 dendritic segments from 14 neurons from 2 cultures; P < 0.001). (**D_3_**), Cumulative distributions of spine length-to-width ratio (shNC: n = 674 spines; sh2#: n = 392 spines; P > 0.999, Kolmogorov–Smirnov test). **E,** Miniature excitatory postsynaptic currents (mEPSCs) recorded from neurons expressing shNC or sh2# (shNC: n = 15 neurons from 2 cultures; sh2#: n = 17 neurons from 2 cultures). (**E_1_**), Representative transfected neurons and mEPSC traces. Quantification of mean mEPSC frequency (Freq, **E_2_,** P < 0.001) and amplitude (Amp, **E_3_**, P = 0.076). **F,** Dendritic spine density and morphology analysis in CA1 pyramidal neurons from Ctrl and HET mice. (**F_1_**), Representative hippocampal brain slices with CA1 neurons labelled with Lenti-hSyn-GFP-Tet-off (Lenti-hSyn-GFP). The boxed region denotes the relative position of dendritic segment used for analysis. Scale bar, 100 μm. (**F_2_**), Representative dendritic segments. Scale bar, 2 μm. (**F_3_**), Quantification of dendritic spine density (Ctrl: n = 31 dendritic segments from 16 neurons and 4 mice; HET: n = 40 dendritic segments from 20 neurons and 5 mice; P < 0.001). (**F_4_**), Cumulative distributions of spine length-to-width ratio (Ctrl: n = 780 spines; HET: n = 679 spines; P > 0.999, Kolmogorov–Smirnov test). **G,** mEPSCs recorded from CA1 pyramidal neurons in acute hippocampal slices from Ctrl and HET mice (Ctrl: n = 19 cells from 6 mice; HET: n = 13 cells from 4 mice). (**G_1_**), Representative traces. Quantification of mean mEPSC frequency (**G_2_**, P = 0.002) and amplitude (**G_3_**, P = 0.158). Data are presented as mean ± s.e.m. in B, C, E and G, as violin plots with the central tendency indicated in D2 and F3. Statistical tests: Two-sided Mann–Whitney U test unless otherwise stated. Exact *P* values are reported for comparisons indicated in the figure; complete statistical results, including nonsignificant comparisons, are provided in the Key resource table 4. Data summary.

We first asked whether TAK1 regulates overall neuronal development or specifically affects excitatory synapses. Acute TAK1 knockdown did not alter dendritic arborization (Fig. 2C). In contrast, dendritic spine density was markedly reduced (Fig. 2D_1_, D_2_), with only limited effects on the morphology of remaining spines (Fig. 2D_3_). These findings suggest that TAK1 is dispensable for dendritic development but is required to maintain spine density.

To determine whether these structural abnormalities were accompanied by impaired synaptic function, we recorded miniature excitatory postsynaptic currents (mEPSCs) from cultured hippocampal neurons (Fig. 2E_1_). TAK1 knockdown significantly reduced mEPSC frequency (Fig. 2E_2_) and produced a rightward shift in the cumulative inter-event interval distribution (Fig. S2D_1_), consistent with the reduction in dendritic spine density. In contrast, the mean mEPSC amplitude was not significantly altered (Fig. 2E_3_), although the cumulative distribution was slightly left-shifted (Fig. S2D_2_), indicating that acute loss of TAK1 primarily affects synapse number rather than the postsynaptic strength of remaining synapses.

To determine whether these cellular phenotypes also occur following chronic neuronal TAK1 reduction in vivo, we sparsely labeled hippocampal CA1 pyramidal neurons using lenti-hSyn-GFP and quantified dendritic spines (Fig. 2F_1_). Consistent with the culture findings, HET mice exhibited a significant reduction in dendritic spine density while largely preserving spine morphology (Fig. 2F). Whole-cell recordings from CA1 pyramidal neurons similarly revealed a significant reduction in mEPSC frequency without detectable changes in mEPSC amplitude (Fig. 2G; Fig. S2E). The milder phenotype observed in vivo is consistent with the partial reduction of TAK1 in HET mice compared with the greater depletion achieved by shRNA-mediated knockdown in cultured neurons.

Together, these findings indicate that neuronal TAK1 supports excitatory synapse structure and function. The concurrent reduction in dendritic spine density and excitatory synaptic transmission further suggested that TAK1 may act locally within excitatory synapses, prompting us to investigate its subcellular localization.

### Postsynaptic localization of TAK1 and its kinase activity support excitatory synapses

Because neuronal TAK1 deficiency impaired excitatory synaptic transmission and dendritic spine structure, we next asked whether TAK1 functions within the postsynaptic compartment. Biochemical fractionation of adult hippocampi revealed substantial expression of TAK1 in both synaptic plasma membrane (SPM) and postsynaptic density (PSD) fractions, together with the respective compartment-specific markers synaptophysin and PSD95 (Fig. 3A; Methods). Independent immunostaining of both exogenously expressed and endogenous TAK1 further revealed punctate TAK1 signals overlapping with PSD95 within dendritic spines (Fig. 3B). These findings indicate that a subset of neuronal TAK1 is present in excitatory postsynaptic compartments.

**Figure 3.**
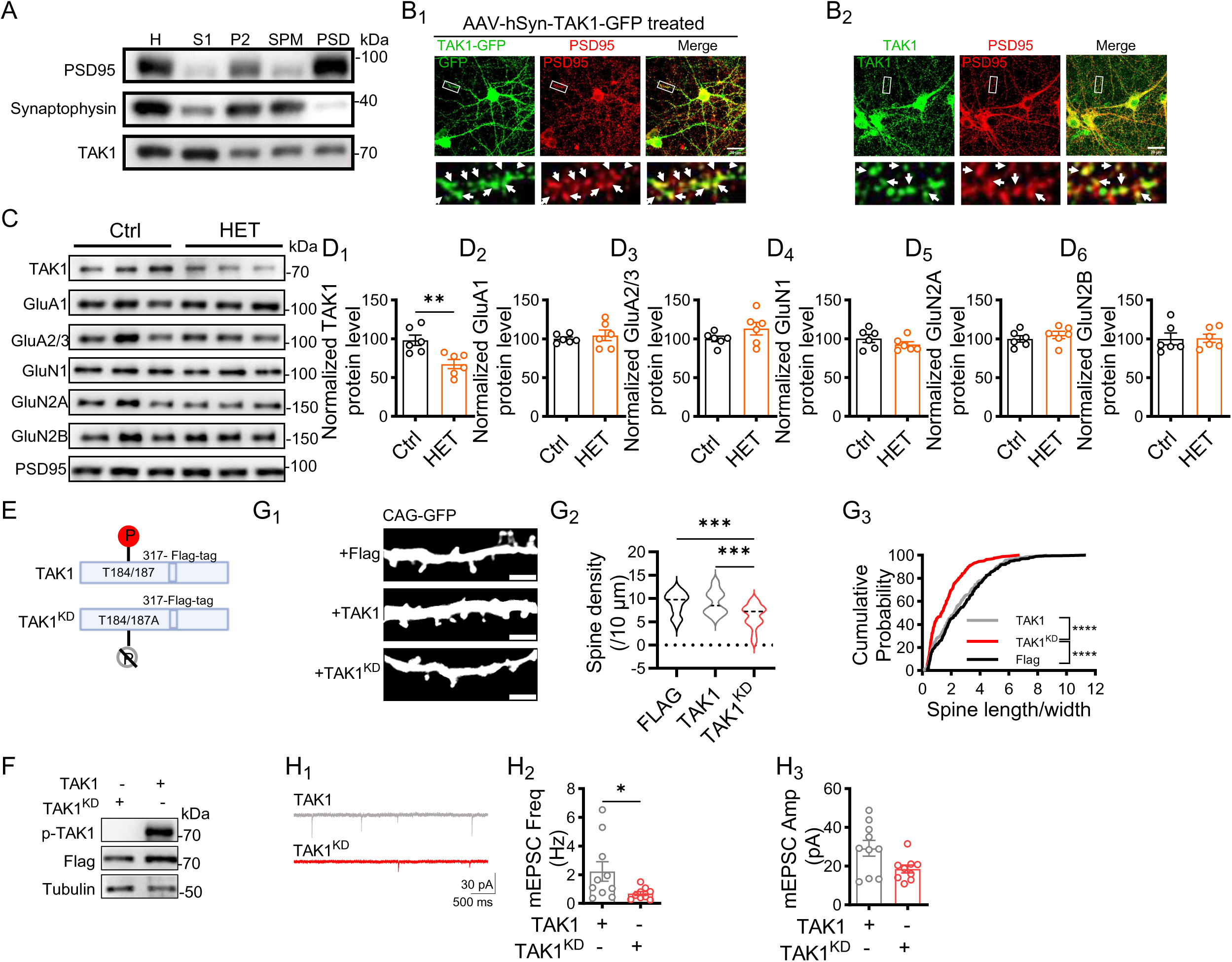
Postsynaptic localization and kinase activity of TAK1 are required for maintaining spine structure and function. **A**, Immunoblot analysis of TAK1, PSD95 and synaptophysin in hippocampal homogenate (H), soluble fraction (S1), crude synaptosomal pellet (P2), synaptic plasma membrane (SPM) and postsynaptic density (PSD) fractions. Molecular masses are indicated in kilodaltons on the right side. **B**, Localization of exogenous and endogenous TAK1 in primary cultured neurons. (**B_1_**), Representative images of neurons expressing AAV2/9-hSyn-3xFlag-eGFP-mMap3k7-WPRE-pA, immunostained for TAK1-GFP (green) and PSD95 (red). (**B_2_**), Representative images of endogenous TAK1 (green) and PSD95 (red). Boxed dendritic regions are enlarged below; arrowheads indicate TAK1 puncta colocalizing with PSD95-positive puncta. Scale bars, 20 μm; enlarged images, 5 μm. **C**, Representative immunoblots of TAK1, GluA1, GluA2/3, GluN1, GluN2A, GluN2B and PSD95 in hippocampal PSD fractions from Ctrl and HET mice. **D**, Quantification of PSD-associated TAK1 (**D_1_**, P = 0.004), GluA1 (**D_2_**, P = 0.725), GluA2/3 (**D_3_**, P = 0.221), GluN1 (**D_4_**, P = 0.333), GluN2A (**D_5_**, P = 0.513) and GluN2B (**D_6_**, P = 0.558), normalized to PSD95 and expressed relative to Ctrl (Ctrl: n = 6 mice; HET: n = 6 mice). **E**, Schematic of wild-type TAK1 carrying a FLAG tag after residue 317 and kinase-deficient TAK1 (TAK1^KD^) carrying alanine substitutions at Thr184 and Thr187. **F**, Western blot validation of TAK1 and TAK1^KD^ expression and TAK1 phosphorylation using tubulin as a loading control. **G**, Effects of TAK1 kinase activity on dendritic spine density and morphology. (**G_1_**), Representative dendritic segments from neurons co-expressing CAG-gag-GFP (CAG-GFP) with Flag, TAK1 or TAK1^KD^. Scale bars, 2 μm. (**G_2_**), Quantification of dendritic spine density (n = 21-24 dendritic segments from 11-12 neurons and 2 cultures; F (2, 64) = 10.17, P < 0.001; P_TAK1 vs. TAK1^KD^ < 0.001; P_Flag vs. TAK1^KD^ < 0.001; One-way ANOVA with Tukey Post Hoc analysis). (**G_3_**), Cumulative distributions of spine length-to-width ratio (Flag_n = 393 spines; TAK1_n = 360 spines; TAK1^KD^_n = 237 spines; P_ TAK1 vs. TAK1^KD^ < 0.001; P_Flag vs. TAK1^KD^ < 0.001, Kolmogorov– Smirnov test). **H**, mEPSCs recorded from neurons co-expressing Lenti-hSyn-GFP with TAK1 or TAK1^KD^ (n = 9-10 neurons from 1 culture). (**H_1_**), Representative traces. Quantification of mean mEPSC frequency (**H_2_**, P_TAK1 vs. TAK1^KD^ = 0.044) and amplitude (**H_3_**, P_TAK1 vs. TAK1^KD^ = 0.113). Data are presented as mean ± s.e.m. in D, H2 and H3 and as violin plots in G2. Statistical tests: Two-sided Mann–Whitney U test was used unless otherwise stated. Exact P values are reported for comparisons indicated in the figure; complete statistical results, including nonsignificant comparisons, are provided in the Key resource table 4. Data summary.

PSD95 organizes postsynaptic signaling complexes that include NMDA receptors. We therefore asked whether TAK1 associates with PSD95. Reciprocal co-immunoprecipitation experiments in both HEK293T and HT22 cells expressing tagged TAK1 and PSD95 showed that the two proteins co-precipitated (Fig. S3A_1_, A_2_). This association was further supported by co-immunoprecipitation of endogenous TAK1 and PSD95 from mouse brain tissue (Fig. S3A_3_). These findings suggest that TAK1 can associate with PSD95-containing postsynaptic protein complexes, providing a potential basis for its postsynaptic localization.

We next examined whether postsynaptic TAK1 was altered in HET mice. Although TAK1 abundance in the soluble hippocampal fraction remained largely unchanged (Fig. S3B, C_1_), PSD-associated TAK1 was significantly reduced (Fig. 3C, D_1_), indicating that partial neuronal TAK1 depletion preferentially affects the postsynaptic pool of TAK1. To determine whether this reduction was accompanied by altered synaptic glutamate receptor abundance, we quantified major AMPA and NMDA receptor subunits in PSD and soluble fractions. Neither PSD-associated nor soluble-fraction receptors levels were significantly altered, except for a modest increase in soluble GluN2B (Fig. 3C, D; Fig. S3B, C). Thus, reduced postsynaptic TAK1 was not accompanied by broad changes in glutamate receptor abundance.

We then asked whether TAK1 kinase activity contributes to excitatory synapse maintenance. Expression of a kinase-dead TAK1 carrying a FLAG tag after residue 317 (TAK1^KD^) mutant, but not wild-type TAK1 carrying a FLAG tag after residue 317 (Fig. 3E, F), significantly reduced dendritic spine density (Fig. 3G_1_, G_2_) and altered morphology of remaining spines (Fig. 3G_3_) without significantly affecting dendritic arborization (Fig. S3D, E). Pharmacological inhibition of TAK1 with 5Z-7-oxozeaenol similarly reduced spine density (Fig. S3F) while leaving dendritic architecture largely intact (Fig. S3G). Consistent with these structural effects, whole-cell recordings showed that TAK1^KD^ expression reduced mEPSC frequency (Fig. 3H_1_, H_2_; Fig. S3H_1_), accompanied by a trend toward reduced mEPSC amplitude (Fig. 3H_3_; Fig. S3H_2_) compared with neurons expressing wild-type TAK1.

Together, these results demonstrate that neuronal TAK1 is present in PSD95-containing postsynaptic compartments and that TAK1 kinase activity supports the structural and functional integrity of excitatory synapses. These findings prompted us to investigate the downstream synaptic substrates through which TAK1 exerts this regulation.

### Postsynaptic phosphoproteomics implicates GluN2B-S920 in TAK1-dependent synaptic regulation

To explore phosphorylation events potentially downstream of postsynaptic TAK1, we performed exploratory quantitative phosphoproteomic profiling of hippocampal postsynaptic density (PSD) fractions from HET and control mice (Fig. 4A). Proteomic and phosphoproteomic measurements were obtained from each individual sample. Principal-component analysis showed separation of HET and control samples in the phosphoproteomic dataset, whereas the corresponding proteomic profiles largely overlapped (Fig. S4A). To distinguish changes in phosphorylation from alterations in protein abundance, phosphosite intensities were normalized to protein levels measured from matched PSD proteomic datasets generated from the same samples (Methods). Fewer than 3% of detected phosphosites were nominally altered in HET mice, approximately 80% of which showed reduced phosphorylation (Fig. 4B; Fig. S4B). This directional pattern is consistent with reduced postsynaptic kinase signaling following partial neuronal TAK1 depletion.

**Figure 4.**
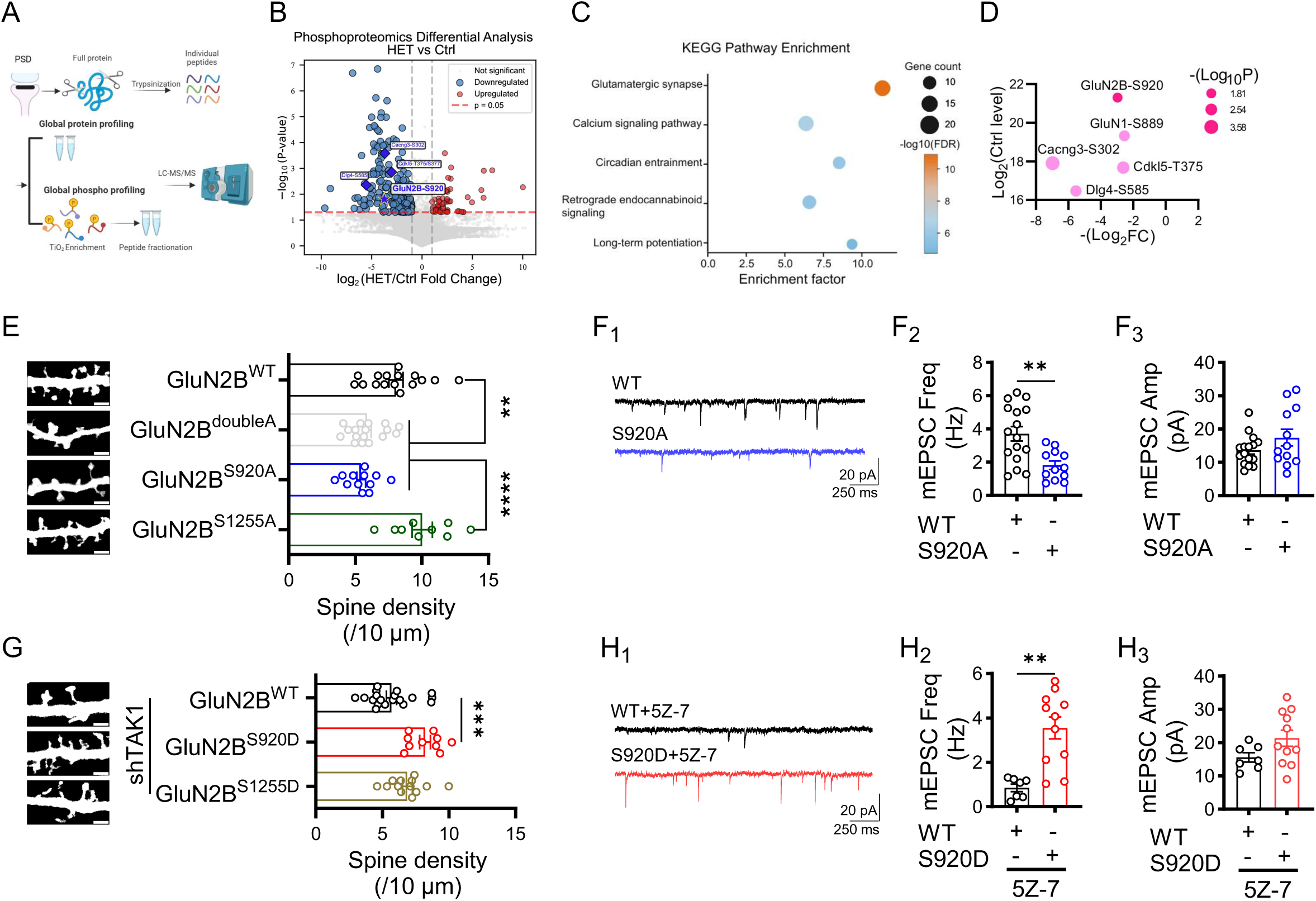
Exploratory phosphoproteomic analysis identifies GluN2B Ser920 as a candidate downstream effector of neuronal TAK1. **A**, Experimental workflow for paired global proteomic and phosphoproteomic analysis of hippocampal PSD fractions from Ctrl and HET mice. **B**, Volcano plot of nominally altered phosphosites in HET relative to Ctrl PSD fractions. The horizontal and vertical dashed lines denote P = 0.05 and |Log_2_(FC)| or |Log_10_(FC)| = 1. Selected synaptic phosphosites, including GluN2B Ser920 (GluN2B-pS920), are labelled (Ctrl: n = 3 biological replicates; HET: n = 4 biological replicates). **C**, KEGG pathway enrichment analysis of proteins containing downregulated phosphosites. The x-axis represents the enrichment factor, circle size and color represent Gene count and −log_10_(FDR-corrected P value). **D**, Selected top-ranked downregulated synaptic phosphosites plotted by fold change and abundance in Ctrl samples. Circle size represents −log_10_(P). **E**, Representative confocal images (left, Scale bar, 2 μm) and quantification (right) of dendritic spine density in primary cultured neurons co-expressing CAG-GFP and GluN2B of different forms (wild-type GluN2B, GluN2B^WT^, n = 16 dendritic segments from 8 neurons and 2 cultures; S920/1255A double mutant GluN2B^doubleA^, n = 17 dendritic segments from 8 neurons and 2 cultures; GluN2B^S920A^, n = 12 dendritic segments from 6 neurons and 2 cultures; GluN2B^S1255A^, n = 9 dendritic segments from 5 neurons and 2 cultures; F (3, 50) = 16.53, P < 0.001; P_WT vs. doubleA = 0.004; P_WT vs. S920A = 0.002; P_doubleA vs. S1255A < 0.001; P_WT vs. S1255A = 0.042; P_920A vs. S1255A < 0.001; One-way ANOVA with Tukey Post Hoc analysis). **F**, mEPSCs recorded from neurons co-expressing Lenti-hSyn-GFP and GluN2B^WT^ (n = 16 neurons from 2 cultures) or GluN2B^S920A^ (n = 12 neurons from 2 cultures). (**F_1_**), Representative traces. Quantification of mean mEPSC frequency (**F_2_**, P = 0.004) and amplitude (**F_3_**, P = 0.397). **G**, Representative confocal images (left, Scale bar, 2 μm) and quantification (right) of dendritic spine density in TAK1-knockdown neurons co-expressing CAG-GFP and GluN2B^WT^, phosphomimetic GluN2B^S920D^ or GluN2B^S1255D^ (GluN2B^WT^_n = 19 dendritic segments from 10 neurons and 2 cultures; GluN2B ^S920D^ _n = 10 dendritic segments from 5 neurons and 2 cultures; GluN2B ^S1255D^ _n = 13 dendritic segments from 7 neurons and 2 cultures; F (2, 39) = 9.414, P < 0.001; P_WT vs. S920D < 0.001; One-way ANOVA with Tukey Post Hoc analysis). **H**, mEPSCs recorded after pharmacological TAK1 inhibition with 5Z-7-oxozeaenol (5Z-7) in neurons co-expressing Lenti-hSyn-GFP and GluN2B^WT^ (n = 7 neurons from 2 cultures) or GluN2B^S920D^ (n = 11 neurons from 2 cultures). (**H_1_**), Representative traces. Quantification of mean mEPSC frequency (**H_2_**, P = 0.003) and amplitude (**H_3_**, P = 0.179). Data are presented as mean ± s.e.m. in F2, F3, H2 and H3. Statistical tests: Two-sided Mann–Whitney U test was used unless otherwise stated. Exact P values are reported for comparisons indicated in the figure; complete statistical results, including nonsignificant comparisons, are provided in the Key resource table 4. Data summary.

Proteins harboring downregulated phosphosites (Fig. S4C) were enriched in pathways related to glutamatergic synapses, calcium signaling, and long-term potentiation (Fig. 4C), suggesting that neuronal TAK1 preferentially influences phosphorylation networks involved in excitatory synaptic function. Several of the downregulated sites with relatively high basal phosphorylation occurred in proteins with established roles in synaptic signaling (Fig. 4D). We therefore assessed the functional consequences of phosphodeficient mutations at selected candidate sites in cultured hippocampal neurons. We first examined two reduced GluN2B phosphosites with high basal phosphorylation in control PSD fractions (Fig. S4D). Expression of the GluN2B-S920A/S1255A double mutant reduced dendritic spine density (Fig. 4E), resembling the effects of TAK1 knockdown and kinase-dead TAK1 expression. Analysis of the individual sites showed that GluN2B-S920A reproduced the reduction in spine density, whereas GluN2B-S1255A had no detectable effect (Fig. 4E). By contrast, phosphodeficient mutants of other prioritized candidates—including GluN1-S889, PSD-95-S585, CDKL5-T375/S377 and CACNG3-S302—did not measurably alter spine density under the same experimental conditions (Fig. S4F–I). Thus, among the candidate phosphosites tested, GluN2B-S920 showed the strongest association with dendritic spine maintenance.

Consistent with its structural effect, expression of GluN2B-S920A significantly reduced mEPSC frequency without affecting amplitude (Fig. 4F; Fig. S4J), closely resembling the synaptic phenotype caused by neuronal TAK1 knockdown or kinase-dead TAK1 expression. We next asked whether a phosphomimetic substitution at S920 could counteract synaptic deficits caused by TAK1 disruption. GluN2B-S920D, but not wild-type GluN2B, increased dendritic spine density in TAK1-knockdown neurons, whereas GluN2B-S1255D produced only a modest, nonsignificant increase (Fig. 4G). Similarly, GluN2B-S920D, but not wild-type GluN2B, counteracted the reduction in excitatory synaptic transmission induced by pharmacological TAK1 inhibition with 5Z-7-oxozeaenol (Fig. 4H and Fig. S4K). Expression of wild-type GluN2B, GluN2B-S920A and GluN2B-S920D in HEK293T cells produced no overt differences in GluN2B or co-expressed GluN1 abundance (Fig. S4E), suggesting that the distinct mutant phenotypes were unlikely to result solely from gross differences in receptor-subunit expression. Together, these findings support a functional contribution of GluN2B-S920 phosphorylation downstream of TAK1 to the maintenance of excitatory synapse structure and function.

### GluN2B-S920 contributes to TAK1-dependent modulation of local synaptic calcium signaling

The enrichment of calcium signaling pathways among the downregulated postsynaptic phosphoproteins in HET mice led us to ask whether TAK1 regulates local calcium dynamics at excitatory synapses. We monitored spontaneous calcium activity at individual dendritic spines in cultured hippocampal neurons using a genetically encoded calcium indicator NEMOf^22^(Fig. 5A). This approach enabled quantification of local calcium transients at synaptic puncta, and somatic calcium activity was assessed separately using Fluo-8 (see Methods).

**Figure 5.**
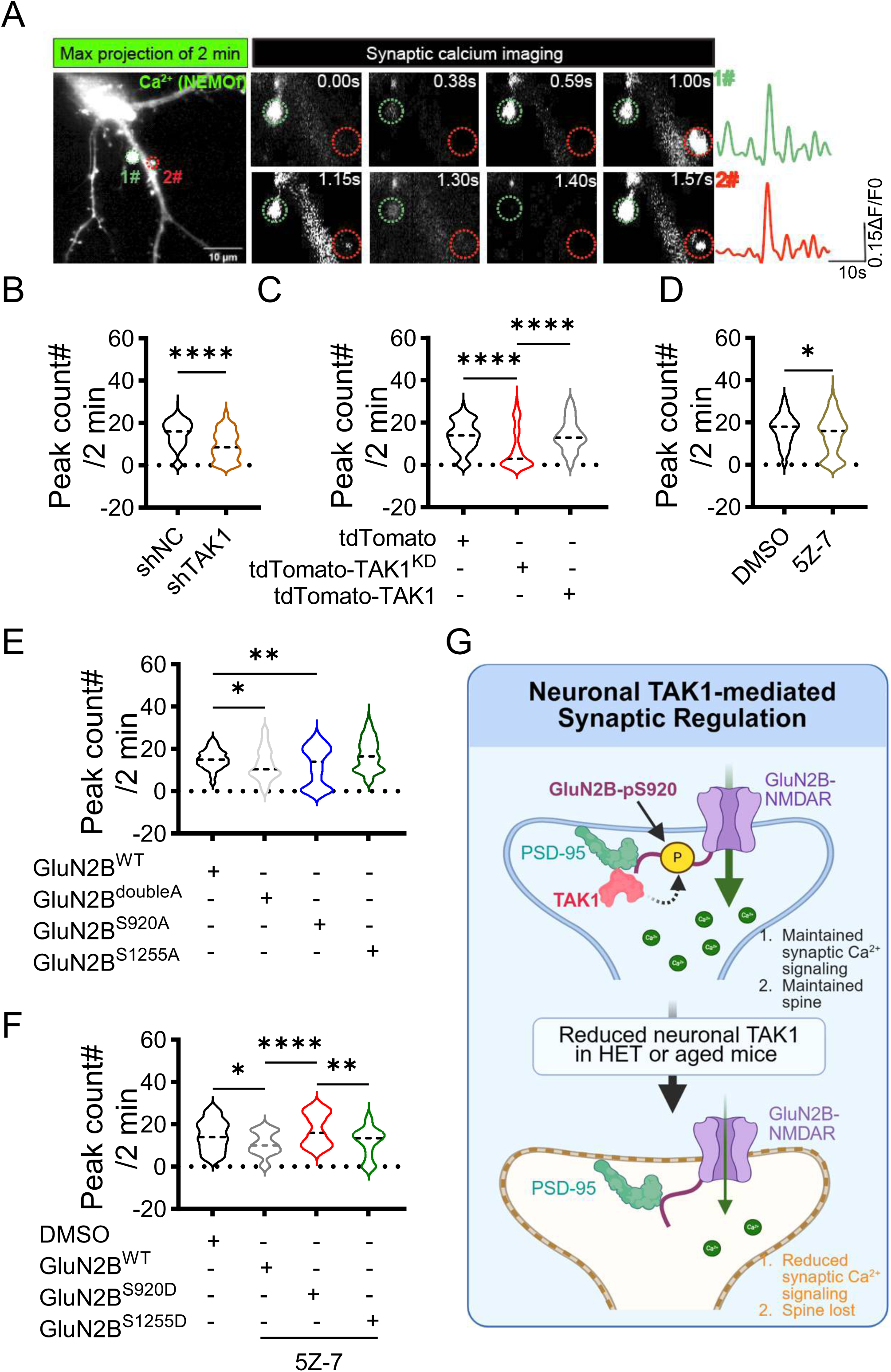
TAK1 and GluN2B Ser920 phosphorylation regulate spontaneous synaptic calcium activity. **A**, Representative synaptic calcium imaging in cultured neurons expressing the calcium indicator NEMOf. The maximum-intensity projection over 2 min, sequential images of two representative synaptic regions and corresponding ΔF/F₀ traces are shown to the right. Scale bar, 10 μm. **B**, Number of synaptic calcium transients detected over 2 min in neurons expressing NEMOf and shNC-mCherry or shTAK1-mCherry (shNC: n = 110 synapses from 11 neurons and 2 cultures; shTAK1: n = 98 synapses from 10 neurons and 2 cultures; P < 0.001). **C**, Number of synaptic calcium transients in tdTomato-expressing control neurons and neurons expressing tdTomato-tagged TAK1^KD^ or wild-type tdTomato-tagged TAK1 (tdTomato_n = 117 synapses from 12 neurons and 2 cultures; tdTomato-TAK1^KD^_n = 100 synapses from 10 neurons and 2 cultures; tdTomato-TAK1 _n = 132 synapses from 14 neurons and 2 cultures; F(3, 346) = 53.56, P < 0.001; P_tdTomato vs. tdTomato-TAK1^KD^ < 0.001; P_tdTomato-TAK1 vs. tdTomato-TAK1^KD^ < 0.001; One-way ANOVA with Tukey’s Post Hoc analysis). **D**, Number of synaptic calcium transients in neurons treated with DMSO or 5Z-7 (DMSO: n = 125 synapses from 13 neurons and 2 cultures; 5Z-7: n = 157 synapses from 16 neurons and 2 cultures; P = 0.027). **E**, Number of synaptic calcium transients in neurons co-expressing NEMOf with tdTomato-tagged GluN2B^WT^, GluN2B^doubleA^, GluN2B^S920A^ or GluN2B^S1255A^ (GluN2B^WT^_n = 125 synapses from 13 neurons and 2 cultures; GluN2B^doubleA^_n = 120 synapses from 12 neurons and 2 cultures; GluN2B^920A^_n = 87 synapses from 9 neurons and 2 cultures; GluN2B^S1255A^_n = 118 synapses from 12 neurons and 2 cultures; F (3, 446) = 12.89, P < 0.001; P_WT vs. doubleA = 0.033; P_WT vs. S920A = 0.007; P_doubleA vs. S1255A < 0.001; P_S920A vs. S1255A < 0.001; One-way ANOVA with Tukey Post Hoc analysis). **F**, Number of synaptic calcium transients after DMSO or 5Z-7 treatment in neurons co-expressing NEMOf with tdTomato-tagged GluN2B^WT^, GluN2B^S920D^ or GluN2B^S1255D^ compared with DMSO (DMSO_n = 37 synapses from 4 neurons and 2 cultures; GluN2B^WT^_n = 51 synapses from 6 neurons and 2 cultures; GluN2B^S920D^_n = 40 synapses from 4 neurons and 2 cultures; GluN2B^S1255D^_n = 30 synapses from 3 neurons and 2 cultures; F (3, 154) = 9.308, P < 0.001; P_DMSO vs. WT = 0.039; P_WT vs. S920D < 0.001; P_S920D vs. S1255D = 0.002; One-way ANOVA with Tukey Post Hoc analysis). **G,** Working model illustrating neuronal TAK1-mediated regulation of excitatory synaptic function and structure. Neuronal TAK1 localized at the excitatory postsynapse is associated with maintenance of GluN2B-pS920 phosphorylation, thereby maintaining synaptic Ca²⁺ signaling and dendritic spine stability. Reduction of neuronal TAK1 decreases GluN2B phosphorylation, leading to impaired synaptic Ca²⁺ dynamics and spine loss. Violin plots show the distribution and central tendency. Statistical tests: Two-sided Mann–Whitney U test was used unless otherwise stated. Exact P values are reported for comparisons indicated in the figure; complete statistical results, including nonsignificant comparisons, are provided in the Key resource table 4. Data summary.

Acute knockdown of TAK1 significantly reduced the frequency of spontaneous synaptic calcium transients (Fig. 5B), whereas somatic calcium activity remained unchanged (Fig. S5A). Expression of kinase-dead tdTomato-TAK1, but not wild-type tdTomato-TAK1, similarly reduced synaptic calcium-event frequency (Fig. 5C), whereas neither construct detectably affected somatic calcium activity (Fig. S5B). Acute pharmacological inhibition of TAK1 with 5Z-7-oxozeaenol also reduced spontaneous synaptic calcium events (Fig. 5D). Thus, genetic and pharmacological disruption of TAK1 converged on a reduction in local synaptic calcium activity, supporting a requirement for TAK1 kinase activity in this phenotype.

We next investigated the source of the TAK1-regulated calcium signals. Inhibition of L-type voltage-gated calcium channels with nifedipine had no detectable effect on event frequency (Fig. S5C). By contrast, two selective antagonists of GluN2B-containing NMDARs, ifenprodil and Ro 25-6981, each reduced spontaneous synaptic calcium events (Fig. S5D, E). These pharmacological results indicate that GluN2B-containing NMDAR activity contributes substantially to the spontaneous local calcium events measured under these conditions.

Given the critical role of GluN2B-S920 in maintaining dendritic spine density, we next asked whether this site also influenced local synaptic calcium signaling. Expression of the phosphodeficient GluN2B-S920A/S1255A double mutant reduced spontaneous calcium-event frequency relative to wild-type GluN2B (Fig. 5E). Analysis of the individual substitutions showed that S920A reproduced this reduction, whereas S1255A had no detectable effect. Thus, among the two TAK1-sensitive GluN2B sites examined, S920 showed the stronger functional association with local synaptic calcium activity.

Finally, we asked whether GluN2B-S920D could counteract the reduction in synaptic calcium activity associated with TAK1 inhibition. In the presence of 5Z-7-oxozeaenol, expression of the phosphomimetic mutant GluN2B-S920D increased synaptic calcium activity to that of control neurons treated with vehicle, whereas wild-type GluN2B or GluN2B-S1255D did not (Fig. 5F). Together, these findings support a model in which the postsynaptic TAK1-GluN2B-S920 phosphorylation axis supports local excitatory synaptic calcium signaling and thereby contributes to the maintenance of excitatory synaptic structure and function (Fig. 5G).

### Neuronal TAK1 re-expression improves hippocampal synaptic plasticity and memory in aged mice

To determine whether the synaptic distribution of TAK1 changes with age, we compared total and postsynaptic TAK1 abundance in the hippocampi of young and aged mice. Total hippocampal TAK1 protein did not differ between the ages examined (Fig. 6A). In contrast, TAK1 abundance in the PSD fraction was significantly reduced in aged mice (Fig. 6B), indicating that aging is associated with a selective reduction of PSD-associated TAK1.

**Figure 6.**
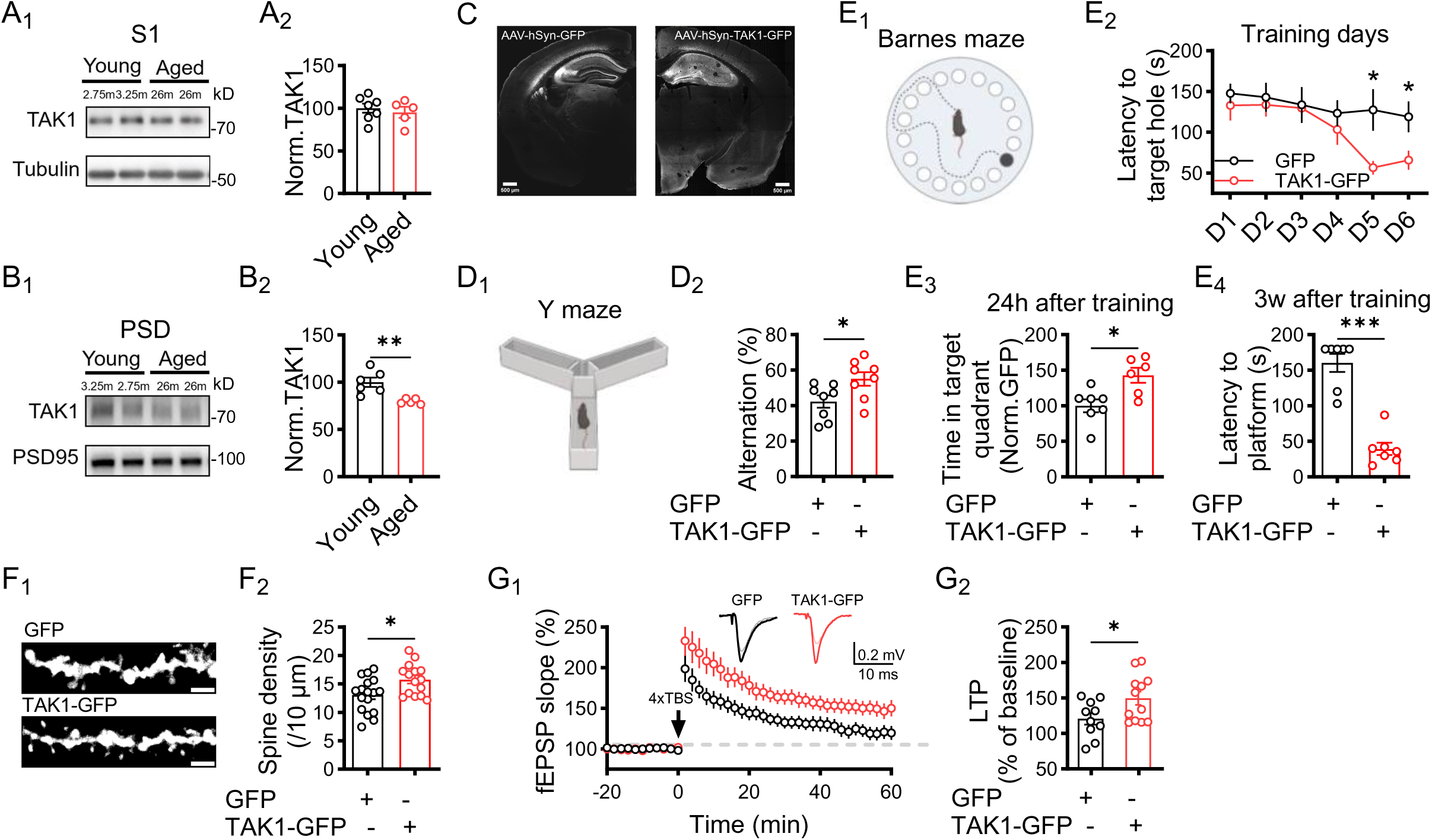
Restoring neuronal TAK1 in the aged hippocampus improves synaptic integrity, plasticity and memory. **A**, TAK1 abundance in hippocampal soluble fractions from young and aged mice. (**A_1_**), Representative immunoblots. (**A_2_**), Quantification of TAK1 normalized to tubulin and expressed relative to young mice (young: n = 7 mice; aged: n = 5 mice; P = 0.639). **B**, TAK1 abundance in hippocampal PSD fractions from young and aged mice. (**B_1_**), Representative immunoblots. PSD95 served as a postsynaptic loading control. (**B_2_**), Quantification of TAK1 normalized to PSD95 and expressed relative to young mice (young: n = 6 mice; aged: n = 5 mice; P = 0.004). **C**, Representative hippocampal sections showing expression of AAV2/9-hSyn-EGFP-WPRE-pA (GFP) or AAV2/9-hSyn-3xFlag-eGFP-mMap3k7-WPRE-pA (TAK1-GFP) in aged mice. Scale bars, 500 μm. **D**, Y-maze performance after hippocampal expression of GFP or TAK1-GFP in aged mice. (**D_1_**), Assay diagram. (**D_2_**), Spontaneous alternation performance (GFP: n = 8 mice; TAK1-GFP: n = 8 mice; P = 0.038). **E**, Barnes maze performance in aged mice expressing GFP or TAK1-GFP. (**E_1_**), Assay diagram. (**E_2_**), Latency to target hole across acquisition training (GFP: n = 7 mice; TAK1-GFP: n = 8 mice; P_day 5 = 0.032; P_day 6 = 0.037, repeated two-sided t-test with Welch correction). (**E_3_**), Time spent in the target quadrant during the probe test 24 h after training, normalized to the GFP group (GFP: n = 7 mice; TAK1-GFP: n = 6 mice; P = 0.022). (**E_4_**), Latency to target hole during the retention test 3 weeks after training (GFP: n = 7 mice; TAK1-GFP: n = 7 mice; P < 0.001). **F**, Dendritic spine density in hippocampal neurons expressing GFP or TAK1-GFP. (**F_1_**), Representative dendritic segments. Scale bars, 2 μm. (**F_2_**), Quantification of dendritic spine density (GFP: n = 17 dendritic segments from 2 mice; TAK1-GFP: n = 15 dendritic segments from 2 mice; P = 0.030). **G**, TBS-induced LTP at Schaffer collateral–CA1 synapses in aged mice expressing GFP or TAK1-GFP. (**G_1_**), Full LTP time course. Insets show representative fEPSP waveforms before (gray and pink) and 60 min after induction (black and red). (**G_2_**), LTP magnitude quantified during the final 10 min of recording (GFP: n = 10 slices from 4 mice; TAK1-GFP: n = 12 slices from 4 mice; two-sided unpaired Welch’s t-test, P = 0.032). Data are presented as mean ± s.e.m. Statistical tests: Two-sided Mann–Whitney U test was used unless otherwise stated. Exact P values are reported for comparisons indicated in the figure; complete statistical results, including nonsignificant comparisons, are provided in the Key resource table 4. Data summary.

We next asked whether increasing neuronal TAK1 expression could improve hippocampal function in aged mice. Bilateral delivery of AAV-hSyn-TAK1-GFP produced neuronal TAK1 expression throughout the hippocampal formation (Fig. 6C). Cpmpared with AAV-hSyn-GFP controls, *Tnf* showed a nonsignificant upward trend, whereas *Il6* expression remained unchanged (Fig. S6A, B).

We then assessed hippocampus-dependent learning and memory. In the Y-maze, aged mice expressing TAK1 exhibited increased spontaneous alternation (Fig. 6D) without altered general locomotor activity (Fig. S6C), consistent with improved spatial working memory. In the Barnes maze, neuronal TAK1 expression improved acquisition during the later training sessions (Fig. 6E_2_), although search strategy distribution was not improved (Fig. S6D). TAK1 -expressing mice spent significantly more time in the target quadrant 24 h after training (Fig. 6E_3_). When retested three weeks later, TAK1-expressing aged mice rapidly located the former target location, whereas GFP controls showed prolonged search latencies, indicating improved long-term memory retention (Fig. 6E_4_). These results demonstrate that re-expressing neuronal TAK1 improved multiple measures of hippocampus-dependent memory in aged mice.

Given the postsynaptic function of TAK1 identified in young adult mice, we next examined whether the cognitive improvement was accompanied by changes in synaptic structure and plasticity. Neuronal TAK1 expression produced a modest but significant increase in dendritic spine density in CA1 pyramidal neurons (Fig. 6F). Theta-burst stimulation-induced long-term potentiation was also enhanced (Fig. 6G), whereas basal excitatory synaptic transmission and paired-pulse ratio remained unchanged (Fig. S6E, F). These findings suggest that neuronal TAK1 supplementation preferentially supports postsynaptic structure and long-term plasticity in the aged hippocampus, consistent with the synaptic role of TAK1 defined in young adult mice.

Together, these results show that aging is accompanied by reduced PSD-associated TAK1 in the hippocampus and that re-expression of neuronal TAK1 improves memory performance, dendritic spine density, and synaptic plasticity in aged mice. These findings support the possibility that reduced postsynaptic TAK1 signaling contributes to age-related synaptic vulnerability.

## Discussion

Here we identify neuronal TAK1 as a previously unrecognized regulator of excitatory synapse integrity and define a postsynaptic TAK1–GluN2B signaling pathway that supports hippocampal plasticity and memory. Partial reduction of TAK1 in forebrain excitatory neurons decreased dendritic spine density and excitatory synaptic connectivity, impaired hippocampal LTP and compromised spatial learning and memory. Mechanistically, a fraction of neuronal TAK1 localized to the postsynaptic density, associated with PSD95-containing complexes and was involved in the regulation of synaptic phosphorylation. Functional analysis identified GluN2B-S920 as an important downstream effector linking TAK1 kinase activity to local synaptic calcium signaling and spine maintenance. Finally, PSD-associated TAK1 was selectively reduced in the aged hippocampus, whereas expression of neuronal TAK1 enhanced spine density, LTP and memory in aged mice. Together, these findings establish postsynaptic TAK1 as a regulator of excitatory synapse integrity and implicate its age-associated reduction in hippocampal synaptic dysfunction.

TAK1 has been studied predominantly as a signaling hub for inflammatory, innate immune and cellular stress pathways, which has likely obscured its physiological importance in neurons^14,15^. Large-scale single-cell atlases show that *Tak1* is broadly expressed across neuronal and non-neuronal populations in the mouse and human brain, including cortical and hippocampal excitatory neurons^19,23,24^. Consistent with these transcriptomic data, our immunohistochemical analyses indicate prominent TAK1 protein expression in neurons of the human cortex and hippocampus (unpublished data). Supporting a neuron-intrinsic function, Harris and colleagues identified *Drosophila* Tak1 as an essential mediator of presynaptic homeostatic plasticity^21^. Our findings extend this concept to mammalian central synapses and reveal a prominent postsynaptic role in maintaining excitatory synaptic structure and signaling. Whereas many well-characterized synaptic kinases are principally discussed in the context of rapid activity-dependent changes in synaptic strength^4,25^, TAK1 appears to support the basal integrity and plasticity competence of mature synapses. This distinction positions the abundance of synaptic TAK1 as a potential determinant of synaptic resilience during aging, a hypothesis worthy of further investigation.

The postsynaptic pool nevertheless represents only a fraction of neuronal TAK1, as TAK1 was most abundant in soluble hippocampal fractions. Its functions elsewhere in the neuron remain unresolved. The reduced paired-pulse ratio in HET mice suggests an additional alteration in presynaptic release probability, consistent with the presynaptic function of TAK1 reported in *Drosophila*^21^. The current data do not determine whether this phenotype reflects loss of TAK1 within axon terminals, developmental adaptation or trans-synaptic compensation. Nevertheless, the combined findings raise the possibility that neuronal TAK1 performs distinct functions according to its subcellular localization: regulating vesicle availability at presynaptic terminals while maintaining phosphorylation-dependent organization at the postsynapse. Whether cytosolic neuronal TAK1 also retains its canonical ability to couple inflammatory or stress signals to downstream pathways remains an important question. The *Drosophila* findings further suggest that neurons may repurpose an evolutionarily conserved innate immune pathway to sense perturbations in synaptic function and initiate compensatory plasticity^21^. Establishing whether mammalian TAK1 similarly integrates immune signals with neuronal activity will require approaches that separate its local synaptic functions from its broader stress-responsive signaling.

Our results provide an initial framework for understanding how TAK1 is positioned at excitatory postsynapses. Reciprocal co-immunoprecipitation supports an association between TAK1 and PSD95, complementing the imaging and biochemical evidence that places TAK1 in the postsynaptic density. PSD95 is a central scaffold that organizes receptors, signaling enzymes and cytoskeletal regulators in excitatory synapses, and the C terminus of GluN2B binds directly to PSD95 PDZ domains^26^. Association with PSD95 could therefore recruit or retain TAK1 within GluN2B-containing signaling complexes, increasing its proximity to GluN2B and other synaptic substrates. Such spatial organization provides a plausible mechanism through which a broadly distributed kinase could achieve local substrate selectivity. Whether there is direct binding or PSD95 is indeed required for synaptic TAK1 localization remains to be examined.

The exploratory phosphoproteomic results imply that TAK1 participates in a broader postsynaptic phosphorylation network rather than regulating a single substrate. Although fewer than 3% of detected sites were altered after partial neuronal TAK1 reduction, most of these sites were downregulated and mapped preferentially to proteins involved in glutamatergic signaling, calcium regulation and LTP. Postsynaptic phosphorylation is organized through interconnected networks of kinases, phosphatases and scaffold proteins, such that perturbation of one kinase can propagate through multiple signaling layers^27,28^. The observed changes are therefore likely to include both direct TAK1 substrates and secondary events arising from altered activity or localization of other synaptic enzymes. Changes in phosphorylation of kinases such as CaMKII and CDKL5 could, for example, influence additional downstream substrates^25,29,30^.

Within this network, GluN2B-S920 emerged as a particularly important functional mediator. GluN2B-S920A reproduced the effects of TAK1 loss on dendritic spine density, mEPSC frequency and spontaneous synaptic calcium activity, whereas GluN2B-S1255A and phosphodeficient mutants of several other highly ranked candidates did not. Conversely, GluN2B-S920D bypassed TAK1 deficiency or inhibition across structural and functional assays. These findings place GluN2B-S920 phosphorylation downstream of TAK1. However, our current data do not establish that TAK1 directly phosphorylates this site. The presence of TAK1 in PSD95-organized NMDAR complexes makes direct phosphorylation plausible, although changes in intermediary kinase or phosphatase activity also remain possible.

The calcium-imaging data provide a functional explanation for the potent synaptic effect of S920. Genetic or pharmacological suppression of TAK1 reduced spontaneous calcium-event frequency at dendritic spines without detectably altering somatic calcium activity. These local events were sensitive to GluN2B-selective antagonists but not to L-type calcium-channel blockade, supporting a major contribution of GluN2B-containing NMDARs. Synaptic NMDAR-mediated calcium signals activate pathways controlling actin remodeling, receptor trafficking and the induction and stabilization of LTP^4,10,31^. Reduced local calcium signaling could therefore weaken the feedback mechanisms that stabilize active spines, providing a potential link between TAK1 deficiency, reduced spine density and impaired LTP.

S920 is located within the proximal portion of the largely intrinsically disordered GluN2B cytoplasmic tail, which is not resolved in available high-resolution NMDAR structures^32^. Previous biophysical and functional studies indicate that phosphorylation and altered conformational dynamics of NMDAR cytoplasmic domains can modulate receptor gating, calcium permeability and interactions with postsynaptic proteins^33–38^. However, these effects are site-specific, and the mechanism through which S920 phosphorylation regulates GluN2B-containing receptors remains to be further explored.

Aging was associated with a selective reduction of PSD-associated TAK1 despite largely preserved total hippocampal TAK1 at the ages examined. This differs from the reduction in total TAK1 reported in aged human cortex by Xu and colleagues¹^8^, potentially reflecting differences in species, brain region, age or biochemical compartment. In mice, impaired synaptic trafficking, anchoring or retention may precede a detectable decline in total protein. Neuronal aging is accompanied by changes in the microtubule cytoskeleton and transport machinery that could compromise the delivery and turnover of proteins at distal synapses^39,40^. Altered PSD95 organization, TAK1 post-translational modification or local proteostasis could also destabilize its synaptic pool. Determining whether TAK1 transport, turnover or association with PSD95 changes with age will be important for explaining the selective synaptic deficit.

Re-expressing neuronal TAK1 in the aged hippocampus improved memory performance, increased spine density and enhanced LTP without detectably altering basal transmission or paired-pulse ratio. The effects were therefore most evident in postsynaptic structure and long-term plasticity, consistent with the function of neuronal TAK1 identified in young adult mice. These experiments suggest that reduced synaptic TAK1 is not merely a correlate of aging but may contribute functionally to declining synaptic resilience.

Synaptic dysfunction and loss are central features of several neurodegenerative disorders and correlate closely with cognitive decline^41–43^. Xu et al. also reported reduced TAK1 in aged human brain and further loss in ALS tissue, although that study emphasized myeloid TAK1 and neuroinflammation^18^. Our findings raise the possibility that loss of neuronal or postsynaptic TAK1 represents an additional axis of vulnerability in diseases characterized by prominent synaptic pathology. Determining whether synaptic TAK1 and GluN2B-S920 phosphorylation are altered in Alzheimer’s disease, ALS/FTD or other synaptopathies will therefore be important.

In summary, this study reveals a physiological neuronal function for a kinase previously viewed largely through the lens of immunity and cellular stress. We propose that TAK1 has compartment-specific neuronal roles and that its postsynaptic pool associates with PSD95-organized complexes and contributes to local calcium signaling and dendritic spine integrity. Exploratory phosphoproteomic and mutational analyses further nominate GluN2B-S920 as a candidate downstream effector of TAK1-dependent synaptic regulation. The age-associated reduction in PSD-associated TAK1, together with the beneficial effects of neuronal TAK1 re-expression in aged mice, raises the possibility that diminished postsynaptic TAK1 signaling contributes to synaptic vulnerability during aging.

## Author Contributions

Conceptualization: Kai-Wen He. Investigation: Yang Liu and Mao-Qing Huang (lead); Yi-Ci Zhang, Xia Min, Zhuo-Ran Jia, Jia-Yao Chen, Fang-Jiao Zong and Jing Gao (supporting). Data analysis: Yang Liu, Mao-Qing Huang, Yi-Ci Zhang, Xia Min, Zi-Yue Zhou and Kai-Wen He. Synaptic calcium imaging method development: Yang Liu, Yiwen Xu, Tong Wang. Writing: Yang Liu, Mao-Qing Huang, Yi-Ci Zhang, Zhuo-Ran Jia, Zi-Yue Zhou, Kai-Wen He. Funding acquisition: Kai-Wen He. Supervision: Kai-Wen He.

## Acknowledgments

The authors thank the staff members of the animal facility at the National Facility for Protein Science in Shanghai (NFPS), Shanghai Advanced Research Institute, Chinese Academy of Sciences, China, for their excellent support. We thank Dr. Youjun Wang from Beijing Normal University for their generous gift of the NEMOf plasmids. This work was supported by Shanghai Municipal Science and Technology Major Project (Grant No. 2019SHZDZX02), Lingang Laboratory (Grant No. LGADB410101), and partly supported by Shanghai Key Laboratory of Aging Studies (19DZ2260400) to Kai-Wen He.

## STAR★METHODS

### METHODS

#### Animals

All procedures were approved by the Institutional Animal Care and Use Committees at the Interdisciplinary Research Center on Biology and Chemistry (IRCBC), Chinese Academy of Science (20230110002, ECSIOC_2026-20), and were conducted in accordance with institutional and national guidelines. Mice were maintained under a 12-h light–12-h dark cycle (7:00 a.m.: defined as zeitgeber time 0, ZT0), with food and water available ad libitum. Aged C57BL/6 mice (18–20 months old) were obtained from Zhejiang Vital River Laboratory Animal Technology Co., Ltd. (Zhejiang, China) or aged in-house at the IRCBC animal facility. To generate forebrain excitatory neuron-specific conditional heterozygous *Tak1* deletion mice, *Tak1*^fl/fl^ mice^44^, kindly provided by Professor Junying Yuan (Shanghai Institute of Organic Chemistry, Chinese Academy of Sciences, Shanghai), were crossed with *Camk2a*-Cre mice (Shanghai Model Organisms, Shanghai, China). Male *Camk2a*-Cre; *Tak1^fl/+^* mice, hereafter referred to as HET mice, were used as the experimental group. Cre-negative *Tak1*^fl/+^ littermates served as controls (Ctrl). Unless otherwise indicated, experiments involving male HET and Ctrl mice were performed at 2–3 months of age.

#### Behavioral Tests

All behavioral training sessions and tests were carried out between ZT4 and ZT8. Mice were transferred to the behavioral testing room at least 30 min before each assay to acclimate to the testing environment.

Experimental groups were tested in a counterbalanced order, and behavioral scoring and analysis were performed by investigators blinded to genotype or treatment. Unless otherwise stated, apparatuses were cleaned with 75% ethanol and allowed to dry completely between animals to minimize olfactory cues.

##### 1) Open field test

Mice were placed in a 40 × 40 cm open-field chamber located in a light- and sound-controlled box under 30–35 lux illumination and allowed to explore freely for 10 min. Mouse activity was recorded by an overhead camera and analyzed using EthoVision XT (Noldus 11.5) or AnyMaze (7.51). Total distance traveled was used to assess locomotor activity, and time spent in the center zone (20 × 20 cm) was used as an indicator of anxiety-like behavior.

##### 2) Elevated-plus maze test

Mice were tested in an elevated plus maze consisting of two open arms (36 × 5 × 0.3 cm) and two closed arms (36 × 5 × 18 cm), elevated 45 cm above the floor under ∼35 lux illumination. Mice were allowed to explore for 10 min, and open-arm time, arm entries, and distance traveled were analyzed using SMART software (Panlab).

##### 3) Three-chamber social-interaction test

Sociability and social-novelty preference were assessed in a three-chamber apparatus consisting of three interconnected chambers (40 × 20 × 23 cm each). The assay consisted of habituation, sociability, and social novelty phases, each lasting 10 min. During the sociability phase, a stranger mouse was introduced into one side chamber, followed by the addition of a second unfamiliar mouse during the social novelty phase. Time spent in each chamber and near the stimulus cages, together with chamber entries, were quantified using EthoVision XT (Noldus 11.5) or AnyMaze (7.51).

##### 4) Y-maze test

Spontaneous alternation was evaluated using a custom-built Y-maze consisting of three identical arms (30 × 6 × 12 cm) positioned 120° apart. Mice were placed at the distal end of a randomly selected arm with the head oriented toward the center of the Y-maze, and allowed to freely explore all three arms for 10 min under dim illumination (∼35 lux). Distance traveled and spontaneous alternation percentage were quantified using EthoVision XT (Noldus 11.5) or AnyMaze (7.51). Spontaneous alternation was calculated as: (successful alternations)/(total arm entries − 2) × 100.

##### 5) Barnes maze test

Spatial learning and memory were assessed using the Barnes maze. The maze consisted of a circular platform (90 cm diameter) with 20 equally spaced holes (5 cm diameter), one of which was connected to an escape box and designated the target hole. The maze was elevated 60 cm above the floor and surrounded by visual cues. Testing was performed under approximately 800 lux illumination with 90-dB white noise as an aversive stimulus.

Following habituation to the maze and escape box, mice underwent two acquisition trials per day for six consecutive days. During each trial, mice were allowed up to 180 s to locate and enter the escape box. Mice that failed to find the escape box were gently guided to it. A probe trial was conducted on day 7 with the escape box removed to assess spatial-memory retention.

The aged mice were retested 3 weeks after the last training to access the long-term memory retention.

For data analysis, mouse trajectories were recorded using ANY-maze (7.51), and time-stamped x–y coordinates were analyzed using Rtrack (v2.0.3)^45^. Search strategies were classified using the Rtrack random forest classifier into spatial and non-spatial strategies. Strategy usage was compared between groups during the learning phase, and spatial strategy preference was analyzed using Fisher’s exact test. The latency to locate the target hole and time spent in the target quadrant during the probe trial were quantified using ANY-maze.

#### Electrophysiology

##### 1) Acute brain slice preparation

Hippocampal slice electrophysiological recordings were conducted following established protocols^43^. In short, mice were deeply anesthetized using isoflurane (#R510-22-10, RWD), followed by rapid decapitation and whole-brain extraction. Both hippocampi were carefully dissected under ice-cold dissection buffer composed of 212.7 mM sucrose, 5 mM KCl, 1.25 mM NaH₂PO₄, 10 mM MgCl₂, 0.5 mM CaCl₂, 26 mM NaHCO₃, and 10 mM glucose, continuously bubbled with 95% O₂/5% CO₂. Transverse hippocampal 300-μm-thick sections (for whole-cell recordings) or 400-μm thick sections (for field recordings) thickness were obtained using a vibratome (Leica, Germany) in the same ice-cold buffer. The resulting slices were then allowed to recover in standard artificial cerebrospinal fluid (ACSF) comprising 119 mM NaCl, 5 mM KCl, 1.25 mM NaH₂PO₄, 26 mM NaHCO₃, 2.5 mM CaCl₂, and 1.5 mM MgCl₂ at 30°C for 30 min, followed by an additional ≥30 min equilibration period at room temperature prior to recording.

##### 2) Whole-cell recordings

Acute hippocampal slices or coverslips containing cultured primary neurons were transferred to a recording chamber and continuously superfused with oxygenated artificial cerebrospinal fluid (ACSF) containing 1.5 mM MgCl₂ and 2.5 mM CaCl₂ for acute brain slices or 1 mM MgCl₂ and 2 mM CaCl₂ for cultured neurons. Pharmacological agents were added as indicated. CA1 pyramidal neurons in acute slices were identified by somatic location and morphology using infrared differential-interference-contrast optics (Olympus).

Recording pipettes (4–6 MΩ tip resistance) were backfilled with various internal solutions (pH 7.2–7.3, osmolarity 280–295 mOsm) tailored to each experimental condition. Recordings were included only when input resistance remained >100 MΩ and access resistance remained <25 MΩ throughout the experiment. Signals were acquired with a MultiClamp 700B amplifier, low-pass filtered at 2 kHz and digitized at 10 kHz using a Digidata 1550B interface (Molecular Devices). AMPA receptor-mediated miniature excitatory postsynaptic currents (mEPSCs) were recorded as previously described^46^. Briefly, 1 μM tetrodotoxin (TTX), 100 μM DL-2-amino-5-phosphonovalerate (D-APV), and 10 μM gabazine were added to the perfusing ACSF to isolate AMPAR mEPSCs. The internal solution contained 125 mM CsOH, 125 mM gluconic acid, 8 mM KCl, 10 mM HEPES, 1 mM EGTA, 4 mM Mg-ATP, 0.5 mM Na₃GTP, 10 mM Na-phosphocreatine, and 5 mM QX-314. Cells were voltage-clamped at a holding potential of −70 mV. Data were analyzed with Mini Analysis Program (Synaptosoft).

##### 3) Field excitatory postsynaptic potential (fEPSP) Recordings

fEPSPs were recorded as described previously^43^. Briefly, slices were maintained in submerged-type recording chambers and superfused with ACSF at a flow rate of 2 ml/min and maintained at 30 ± 0.5°C. Schaffer collateral fibers were stimulated using 0.2-ms constant-current pulses delivered via custom-built double-barreled glass microelectrodes (Sutter Instrument Co.) filled with ACSF, with stimulus intensities adjusted to yield half-maximal fEPSP amplitudes. Evoked responses were recorded from the proximal CA1 stratum radiatum, bandpass-filtered at 0–1000 Hz, digitized at 10 kHz, and stored using WinLTP software (WinLTP Ltd.). The initial slope of each fEPSP was subsequently measured with WinLTP Reanalysis.

###### a. Paired-pulse ratio (PPR)

Paired stimuli were delivered at inter-stimulus intervals (ISIs) of 25, 50, 100, 250, and 500 ms. The PPR value was determined by dividing the slope of the second response by that of the first.

###### b. Input-Output (I/O) relationship

Input–output relationships were generated by progressively increasing stimulus intensity and recording four or five responses at each intensity. The amplitude of fiber volley was measured and used as the input strength.

###### c. Long-term potentiation (LTP)

Baseline responses were evoked at 0.033 Hz using single-pulse stimulation at an intensity set to 30% of the half-maximal level. Following a stable baseline period of 20–30 min, LTP was induced by four theta-burst stimulation (TBS) trains delivered at 0.1 Hz. Each TBS train comprised 10 bursts, with each burst containing four pulses at 100 Hz, repeated at 5 Hz.

###### d. Long-term depression (LTD)

A 20-min stable baseline was first established using paired-pulse stimulation (ISI = 50 ms) at 0.033 Hz. LTD was subsequently induced by applying a paired-pulse 1 Hz protocol (PP-1 Hz), consisting of paired pulses with a 50-ms ISI delivered continuously at 1 Hz for 15 minutes.

#### Stereotaxic Injection

Mice were anesthetized via a face mask with 1% isoflurane delivered in air at a flow rate of 0.5 mL/min and stereotactically injected with AAV encoding TAK1 or control GFP into the hippocampus. Stereotaxic injections were performed at the following coordinates: AP, −2.70 mm and ML, ±2.25 mm relative to bregma, and DV, −1.35 mm relative to the skull surface. The viral suspension was infused at a rate of 200 nL/min. After injection, the needle was left in place for 10 min before being slowly withdrawn. AAV 2/9-hSyn-EGFP-WPRE-pA and AAV2/9-hSyn-3xFlag-eGFP-mMap3k7-WPRE-pA (Shanghai Taitool Bioscience Co.) were diluted to a titer of 10^11^ vg/mL, and each mouse received a bilateral injection of 1 μL of the viral suspension for each hippocampus. After surgery, the animals were allowed to recover fully from anesthesia before being returned to their home cages. After 28 days, animals were subjected to spine density analysis by immunostaining, electrophysiology studies, or behavior tests before being sacrificed and analyzed to verify the injection accuracy and infection efficiency.

GFP-Tet-off lentiviral vectors (500 nL) were also stereotactically injected into hippocampus of Ctrl and HET mice. 14 days later, the animals were deeply anesthetized and transcardially perfused with ice-cold PBS, followed by 4% paraformaldehyde (PFA) in PBS. Brains were subsequently removed and post-fixed in 4% PFA at 4 °C for overnight and analyzed spine density by immunofluorescence staining with an anti GFP antibody.

#### Primary Neuronal Culture and Transfection or Transduction

Primary mouse hippocampal neurons were prepared from E16–18 C57BL/6J mouse embryos (Shanghai Ling Chang Biotechnology Co.). Primary rat cortical and hippocampal neurons were isolated from P0 Sprague–Dawley rats (Zhejiang Vital River Laboratory Animal Technology Co., Ltd., Zhejiang, China). The isolated neurons were then cultured in Neurobasal medium supplemented with 2% B-27, 0.25% GlutaMAX, and 1% penicillin–streptomycin at 37 °C with 5% CO₂.

Mouse neurons were transduced with AAV or lentivirus, or transfected with plasmids using Lipofectamine 3000 (Thermo Fisher Scientific) following the manufacturer’s instructions at DIV7–8. All downstream experiments were carried out at DIV15-16.

For Lipofection, two-plasmid co-transfection was performed using 0.4 μg of each plasmid and 1 μL of Lipofectamine 3000 reagent per well. For three-plasmid co-transfection, 0.35 μg of each plasmid and 1.2 μL of Lipofectamine 3000 reagent were used per well. For lentiviral or AAV transduction, primary neurons were treated with 1 μL per well. Detailed information on the viral vectors and plasmid constructs is provided in Key resource table 1.

Rat neurons were treated with 5-fluoro-2′-deoxyuridine and uridine to suppress glial proliferation at DIV4. At DIV12, neurons were co-transfected with NEMOf and indicated plasmids (1μg of each plasmid) using 1μL Lipofectamine 3000 according to the manufacturer’s instructions. Synaptic calcium imaging was performed at DIV15–16.

#### Immunofluorescence Staining

For mouse brain samples, including lentivirus-injected Ctrl/HET mice and AAV-injected aged mice, animals were transcardially perfused with ice-cold PBS followed by 4% paraformaldehyde (PFA). Brains were post-fixed in 4% PFA overnight, cryoprotected sequentially in 20% and 30% sucrose for at least 12 h at each concentration, or until they sank to the bottom, embedded in OCT compound, and stored at −80 °C. Next, 25-μm coronal slices were sectioned using a Leica cryostat (model: CM3050s-1-1-1) and mounted on poly-lysine-coated glass slides. Slices were stored at -20°C. Before staining, slices were rinsed with PBS and incubated for 2 h at room temperature with a blocking solution consisting of 5% normal goat serum (Sigma) and 0.1% Triton X-100 in PBS.

For primary neurons, cells cultured on coverslips were washed three times with PBS and fixed with fixation buffer (4% PFA and 4% sucrose in PBS) for 10 min at room temperature. The cells were then permeabilized with PBS containing 0.15% Triton X-100 for 15 min. Next, the cells were incubated with blocking solution (5% bovine serum albumin in PBS) for 30 min at room temperature.

Brain sections or cultured cells were incubated with primary antibodies at 4°C overnight, followed by washing three times with PBST buffer (0.1% Tween 20 in PBS). They were then incubated with secondary antibodies at room temperature for 2 h and DAPI for 5 min in the dark, followed by washing three times with PBST. Next, they were mounted with mounting medium (Invitrogen, ProLong Gold antifade reagent) for imaging.

#### Confocal imaging and morphometric analysis of neurons

Dendritic spine density in CA1 pyramidal neurons was assessed using a Leica TCS SP8 X laser-scanning confocal microscope. Secondary and tertiary basal dendrites were imaged using 488-nm excitation and a 100× oil-immersion objective. Images were acquired as z-stacks with voxel sampling of 0.036 × 0.036 × 0.217 μm in the x, y and z dimensions, respectively. For each neuron, two to four dendritic segments were imaged. Z-stacks were projected and spine density was quantified in ImageJ (NIH) by an investigator blinded to experimental condition.

Dendritic morphology in primary mouse neurons was imaged using an Andor spinning-disk confocal microscope equipped with a 20× air objective. Z-stacks were acquired with voxel sampling of 0.3 × 0.3 × 1 μm in the x, y and z dimensions, respectively. Following z-projection, dendritic arborization was quantified using the Sholl Analysis plugin in ImageJ by an investigator blinded to experimental condition.

Hippocampal AAV or lentiviral injection sites were visualized using a 20× air objective in tile-scan mode. Representative images were acquired to document the anatomical distribution of viral expression.

#### Calcium Imaging and Data Analysis

##### 1) Synaptic calcium imaging

Primary rat neurons were transfected at DIV7 with a plasmid encoding the genetically encoded calcium indicator NEMOf (1 μg/3.5 cm dish). At DIV15–16, the culture medium was replaced with calcium-imaging artificial cerebrospinal fluid (ACSF containing 2.2 mM CaCl₂, 5.6 mM D-glucose, 0 mM Mg²⁺, pH 7.4). Time-lapse images were acquired for 2 min at 37 °C and 5% CO₂ using a structured-illumination microscope (SIM, CSR Biotech) with a 100× oil-immersion objective at 37°C with 5% CO_2_. Images were sampled at 0.065 × 0.065 μm per pixel and acquired every 150 ms.

To assess the channel dependence of spontaneous synaptic calcium activity, neurons were first imaged for 2 min under vehicle conditions to establish baseline activity. Ro 25-6981 (0.5 μM), ifenprodil (3 μM) or nifedipine (8 μM) was then added, and imaging continued for an additional 2 min using identical acquisition settings. Baseline and post-treatment recordings were analyzed as paired measurements from the same neurons, except in the nifedipine group, for which recordings were obtained from different neurons and analyzed as unpaired measurements.

##### 2) Somatic calcium imaging

Primary mouse hippocampal neurons were transduced at DIV8 with respective lentiviruses (shNC/shTAK1/tdTomato/tdTomato-TAK1/tdTomato-TAK1^KD^). At DIV16, neurons were incubated for 1 h with 4 μM Fluo-8 in ACSF. The dye-containing solution was then replaced with calcium-imaging ACSF containing 2.2 mM CaCl₂, 5.6 mM d-glucose and no added Mg²⁺ (pH 7.4).

Time-lapse images were acquired for 2 min at 37 °C with constant 5% CO₂ using an FV4000 microscope equipped with a 20× air objective and 488-nm excitation. Images (0.139 × 0.139 μm per pixel) were acquired every 150 ms.

##### 3) Calcium-signal analysis

Time-lapse images were analyzed in ImageJ and custom Python scripts. Individual dendritic spines or individual neuronal somata were manually defined as regions of interest (ROIs), and fluorescence intensity at each time point was recorded as *F_t_*. The first ten frames (1.5 s) of each recording were excluded to minimize acquisition-related artifacts. Baseline fluorescence (*F*_0_) was defined as the 20^th^ percentile of fluorescence values across the remaining trace. Fluorescence changes were calculated as Δ*F* = *F_t_* − *F*_0_ and expressed as 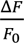. The resulting traces were plotted over time, and calcium-event frequency was quantified using custom Python code.

#### Molecular Biology

##### 1) Construction of Plasmids, Reagents, and Antibodies

AAV9-hSyn-EGFP-WPRE-pA and AAV9-hSyn-3×FLAG-EGFP-mMap3k7-WPRE-pA were custom produced by Shanghai Taitool Bioscience. Viral stocks had an initial titer of 1 × 10^13^ vg/mL and were diluted to 1 × 10^11^ vg/mL for experimental use.

An shRNA targeting mouse Tak1 (5′-TCTGAGAGGAAGGCTTTCATT-3′) was cloned into pCLenti-U6-shRNA-UbC-mCherry-WPRE. A non-targeting shRNA (shNC; 5′-TTCTCCGAACGTGTCACGT-3′) served as the negative control. Both lentiviral constructs were generated by OBiO Technology (Shanghai, China).

The following expression plasmids were constructed using the pRRLsin-PPTs-hCMV lentiviral vector backbone: full-length GluN2B (Gene ID: 14812, GluN2B^WT^), GluN2B^S920A^, GluN2B ^S1255A^, GluN2B-S920A/S1255A (GluN2B^doubleA^), GluN2B^S920D^, GluN2B^S1255D^, full-length GluN1 (Gene ID:14810, GluN1^WT^), GluN1^S889A^,GluN1^S889D^, full-length PSD95 (Gene ID: 13385, PSD95^WT^), PSD95^S585A^,PSD95^S585D^, full-length CDKL5 (Gene ID:382253, Cdkl5^WT^), CDKL5^T375A/S377A^, CDKL5^T375D/S377D^, full-length Cacng3 (Gene ID: 54376, Cacng3^WT^), Cacng3^S302A^, Cacng3^S302D^. The full-length mouse TAK1 construct was kindly provided by the laboratory of Dr. Junying Yuan^47^. For detection, an in-frame FLAG epitope was inserted immediately after amino acid 317 of TAK1. Kinase-dead TAK1 (TAK1^KD^) was generated by substituting alanine for T184 and T187. FLAG-tagged wild-type TAK1 and TAK1-T184A/T187A were cloned into a lentiviral expression backbone. tdTomato–TAK1 and tdTomato–TAK1^KD were generated by fusing tdTomato to the N terminus of the corresponding TAK1 construct. All constructs and introduced mutations were verified by DNA sequencing.

The hSyn-GFP-Tet-Off lentiviral vector (Lenti-hSyn-GFP) was used for dendritic spine labeling in vivo and for whole-cell recordings in cultured neurons, as described previously^43^. The genetically encoded calcium indicator NEMOf was kindly provided by Professor Youjun Wang (Beijing Normal University, Beijing)^48^. The CAG-gag-GFP plasmid used for morphological labeling of cultured neuron was kindly provided by Professor Yelin Chen (Shanghai Institute of Organic Chemistry, Chinese Academy of Sciences, Shanghai).

All reagents and antibodies are listed in Supplementary Key resource table 2 and 3, respectively.

##### 2) Lentivirus production

Lentiviruses were produced as described previously^43^. Briefly, HEK293T cells were transiently transfected with the lentiviral expression construct and the packaging plasmids psPAX2 and pMD2.G using polyethyleneimine (PEI; Titan Scientific) according to the manufacturer’s instructions. Half of the culture medium was replaced 6–8 h after transfection. Viral supernatants were collected 48 h after transfection, cleared of cellular debris and passed through a 0.45-μm filter. Lentiviral particles were concentrated by ultracentrifugation at 20,000 r.p.m. for 2 h at 4 °C. Pellets were resuspended in PBS at 25 μl per 10 ml of starting supernatant, aliquoted and stored at −80 °C until use.

##### 3) RNA extraction

Mice were anesthetized with isoflurane and euthanized. Hippocampi were rapidly dissected, transferred to RNase-free tubes and snap-frozen in liquid nitrogen. Primary mouse neurons were collected at DIV16 and lysed directly in 100 μl TRIzol reagent.

Total RNA was extracted using TRIzol (Invitrogen) and treated with the TURBO DNA-free kit (Invitrogen) according to the manufacturers’ instructions. RNA concentration and purity were assessed using an ND-1000 spectrophotometer (NanoDrop Technologies).

##### 4) Quantitative reverse-transcription PCR

Quantitative reverse-transcription PCR (RT–qPCR) was performed using the QuantiTect SYBR Green RT-PCR kit (Qiagen). Primers targeting *Tak1, Il6, Tnf* and *Actb* were obtained from Sangon Biotech. Actb was used as the reference gene, and all reactions were performed in triplicate. Relative transcript abundance was determined from SYBR Green fluorescence using the comparative threshold-cycle method.

The following qPCR primers were used in this study: *Tak1*-F, 5’-ACCAGGCAAAGCAACAGAGT-3’; *Tak1*-R, 5’-GTTGCCCGTTACCTGCAGTC-3’; *Il6*-F, 5 ′ ACAAAGCCAGAGTCCTTCAGAGA-3′; *Il6*-R, 5′-AGGAGAGCATTGGAAATTGGGGT-3′; *Tnf-α*-F, 5′-ATGAGAAGTTCCCAAATGGC-3’; *Tnf-α*-R,5′-CTCCACTTGGTGGTTTGCTA-3′; *Actb*-F, 5′-GTGTGATGGTGGGAATGGGT-3’; *Actb*-R,5′-GCTGGGGTGTTGAAGGTCTC-3’.

#### Biochemistry

##### 1) PSD preparation and Western Blotting

Postsynaptic density (PSD) preparation was carried out as previously describe^43^. Briefly, brain lysates were prepared by Dounce homogenization (H) and differential centrifugation. Following centrifugation at 1,000 × g for 10 min, the nuclear pellet and debris (P1) were removed, and the supernatant (S1) was centrifuged at 12,000 × g for 20 min to obtain the P2 fraction. Both P1 and P2 fractions were washed twice to remove nonspecific contaminants. The P2 fraction was subjected to Triton X-100 extraction (0.5% Triton X-100, 4°C, 15 min) and centrifuged at 12,000 × g for 20 min to separate detergent-soluble (SPM) and detergent-insoluble fractions (PSD). The insoluble fraction was dissolved in 2% SDS for subsequent analysis.

S1 (10 μg) and PSD (1 μg) proteins were separated on 8% SDS–polyacrylamide gels and transferred to polyvinylidene difluoride membranes (Bio-Rad). Membranes were blocked for 1 h at room temperature in Tris-buffered saline containing 0.1% Tween-20 (TBST) and 5% non-fat milk, followed by incubation with primary antibodies overnight at 4 °C. After washing with TBST, membranes were incubated with the appropriate secondary antibodies for 1 h at room temperature. Immunoreactive bands were detected using a Fusion chemiluminescence imaging system (Vilber). Protein abundance in S1 fractions was normalized to α-tubulin, whereas abundance in PSD fractions was normalized to PSD-95.

##### 2) Co-immunoprecipitation

Co-immunoprecipitation was used to assess the association between TAK1 and PSD-95 in mouse hippocampal tissue, HEK293T cells and HT22 cells. Hippocampi were homogenized in ice-cold 4 mM HEPES buffer supplemented with protease and phosphatase inhibitors. Homogenates were centrifuged at 20,000 × g for 15 min at 4 °C, and the supernatants were collected. Protein concentrations were determined using a BCA assay. For each immunoprecipitation, 1 mg of protein was used, and 40 μl of lysate was retained as the input sample.

For cell-based assays, HEK293T or HT22 cells were transfected with the indicated plasmids. For PSD-95 immunoprecipitation, cells were co-transfected with TAK1 and PSD-95 or with TAK1 and the corresponding empty vector. For reciprocal TAK1 immunoprecipitation, cells were co-transfected with PSD-95 and TAK1 or with PSD-95 and empty vector. Cells were collected 48 h after transfection and lysed in 1 ml of ice-cold lysis buffer supplemented with protease and phosphatase inhibitors. Lysates were centrifuged at 20,000 × g for 15 min at 4 °C, and equal amounts of soluble protein were used for immunoprecipitation.

Lysates were incubated overnight at 4 °C with an antibody against PSD-95 or TAK1, or with the corresponding species-matched control IgG, with continuous rotation. Pre-washed magnetic beads were then added, and samples were incubated for an additional 1 h at 4 °C. Bead– antibody–protein complexes were recovered using a magnetic rack and washed three times with lysis buffer. Bound proteins were eluted in 30 μl of 1× SDS sample buffer by heating at 95 °C for 5 min and analyzed by immunoblotting. Input samples represented approximately 1– 2% of the protein used for immunoprecipitation.

#### Mass Spectrometry–based Phosphoproteomic Analysis

##### 1) Protein digestion and phosphopeptide enrichment

Postsynaptic density (PSD) fractions isolated from mouse hippocampi were precipitated on ice with four volumes of cold acetone. Protein pellets were collected by centrifugation at 14,000 × g for 15 min and dissolved in 8 M urea and 500 mM Tris–HCl, pH 8.5. Proteins were reduced with 5 mM tris(2-carboxyethyl) phosphine for 20 min and alkylated with 10 mM iodoacetamide for 15 min at room temperature in the dark. Samples were diluted to 2 M urea with 100 mM Tris–HCl, pH 8.5, containing 1 mM CaCl₂ and digested overnight at 37 °C with sequencing-grade trypsin (Promega, V5111) at an enzyme-to-protein ratio of 1:100 (w/w). Peptides were desalted using C18 StageTips.

For phosphoproteomic analysis, phosphopeptides were enriched using TiO₂ resin (GL Sciences) packed into C8 StageTips. Peptides were loaded in 1 M glycolic acid, 5% trifluoroacetic acid and 80% acetonitrile. After sequential washing, bound phosphopeptides were eluted, dried under vacuum and reconstituted in 0.1% formic acid for liquid chromatography–tandem mass spectrometry (LC–MS/MS). An unenriched peptide aliquot was analyzed in parallel to quantify total protein abundance.

##### 2) LC–MS/MS acquisition

Peptides were analyzed using a nanoElute liquid chromatography system coupled to a timsTOF Pro 2 mass spectrometer (Bruker). Peptides were separated on a 25-cm C18 analytical column (75-μm inner diameter; 1.5-μm particle size) maintained at 50 °C and operated at 300 nL min⁻¹. Mobile phases were 0.1% formic acid in water (A) and 0.1% formic acid in acetonitrile (B). A 60-min gradient was used for proteomics samples, and a 120-min gradient was used for phosphoproteomics samples. All spectra were acquired within an m/z range of 100 to 1700 and an ion mobility range from 0.75 to 1.40 V.s/cm^2^. The dual TIMS analyzer was operated at a fixed duty cycle with a ramp time of 100 ms, and the total cycle time was 1.17 s. Data-dependent acquisition (DDA) was performed in PASEF mode, with 10 PASEF scans per topN acquisition cycle.

##### 3) Database searching and label-free quantification

Raw files were processed in FragPipe v.20.0. Spectra were searched against the UniProtKB mouse protein database (downloaded in March 2024) supplemented with common contaminants and decoy sequences. Trypsin specificity was applied, allowing up to one missed cleavage. Carbamidomethylation of cysteine was specified as a fixed modification, and methionine oxidation and protein N-terminal acetylation as variable modifications. For phosphoproteomic searches, phosphorylation of serine, threonine and tyrosine (+79.9663 Da) was included as an additional variable modification.

Peptide-spectrum matches, peptides and proteins were controlled at a 1% false-discovery rate using Percolator and ProteinProphet within Philosopher v.5.0.0. Phosphosite localization was evaluated using PTM-Prophet, and sites with a localization probability of at least 0.75 were retained. Label-free peptide and protein abundances were quantified using IonQuant v.1.9.8 with match-between-runs enabled.

##### 4) Phosphoproteomic data processing and differential analysis

Label-free phosphopeptide intensities and corresponding total-protein abundances from control (Ctrl) and conditional heterozygous *Tak1* deletion (HET) samples were analyzed using a custom Python workflow. Non-positive and missing abundance values were treated as missing. Intensities were log2-transformed, after which missing values were imputed separately for each sample with a deterministic low-abundance value defined as the minimum observed log_2_ intensity in that sample minus 1.8 times the standard deviation of its observed log_2_ intensities. If the standard deviation was zero or could not be estimated, a value one log_2_ unit below the observed minimum was used. Phosphoproteomic and total-proteomic matrices were then median-normalized independently by centering each sample to the global median of the corresponding dataset.

Phosphosites were matched to their parent proteins using protein accession identifiers (Protein.Ids). When multiple protein-level entries mapped to the same accession, their log2 abundances were consolidated by the median. For each matched sample, protein-corrected phosphosite abundance was calculated as:

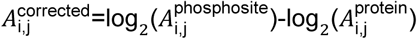

where denotes the phosphosite, its corresponding parent protein and the biological sample. Phosphosite entries without a matched parent-protein measurement were excluded from protein-corrected analyses. Because phosphosite and total-protein LFQ intensities have dataset-specific arbitrary scales, corrected values were interpreted as relative protein-adjusted phosphosite abundances rather than absolute phosphorylation stoichiometries.

Protein-corrected abundances were compared between Ctrl and HET samples using two-sided unpaired Welch’s -tests. Log_2_ fold changes were calculated as the difference between the mean corrected log_2_ abundance in HET and Ctrl samples. Phosphosites with an absolute log_2_ fold change greater than 1 and a nominal P<0.05 were classified as candidate differentially regulated sites. The candidate-selection threshold was based on nominal values because of the limited sample size. Accordingly, the phosphoproteomic analysis was considered exploratory and was used to prioritize candidates for independent experimental validation.

##### 5) Phosphosite annotation and consolidation

Phosphorylated residues were annotated directly from the amino-acid identity and protein-coordinate fields PTM.SiteAA and PTM.SiteLocation, respectively^50^. Protein accessions were obtained from Protein.Ids; where available, gene symbols were extracted from the corresponding protein descriptions. Each site was labeled by gene or protein identifier, modified residue and protein position. Entries mapping to the same annotated phosphosite were consolidated by reporting the median log_2_ fold change and the lowest nominal value across corresponding entries. Serine, threonine and tyrosine sites were summarized separately.

##### 6) Pathway enrichment analysis

Genes corresponding to candidate phosphosites with decreased phosphorylation were subjected to Kyoto Encyclopedia of Genes and Genomes (KEGG) pathway enrichment analysis using a hypergeometric test. Enrichment *P* values were adjusted using the Benjamini– Hochberg procedure, and pathways with an adjusted *Q* < 0.05 were considered significantly enriched. Analyses were performed in Python using [Version 3.14].

#### Statistical Analysis

The corresponding statistical test and number of samples were indicated in the figure and figure legends for each experiment. Statistical analysis was performed by Prism V11.0 software (GraphPad Software, Inc.). Results are presented as mean ± SEM. Mann-Whitney U test was used for unpaired data. Unpaired two-tailed Student’s t-test was used to analyze the phosphoproteomics data. Two-sided Wilcoxon matched-pairs signed rank test was used for paired data. One-way ANOVA followed by post hoc multiple comparison test (Tukey’s) was used to compare data with more than 2 groups. Two-way ANOVAs with Sidak post hoc multiple-comparisons test were used to compare data with more than two groups and two independent variables. Cumulative distributions were compared by Kolmogorov-Smirnov (KS) test. Error bars in all figures indicate standard error of mean (s.e.m). Only significant comparisons were labeled in the figures. The level of significance was set at P < 0.05. *P <0.05; * *P < 0.01; * **P < 0.001; * ** *P < 0.0001.

## Data Resource and code availability

The phosphoproteomic and proteomic raw data and customized Python codes are available on GitHub: https://github.com/KaiWen-Helab/2026Liu-et-al

## Supplementary Figure Legends

**Supplementary Figure 1.**
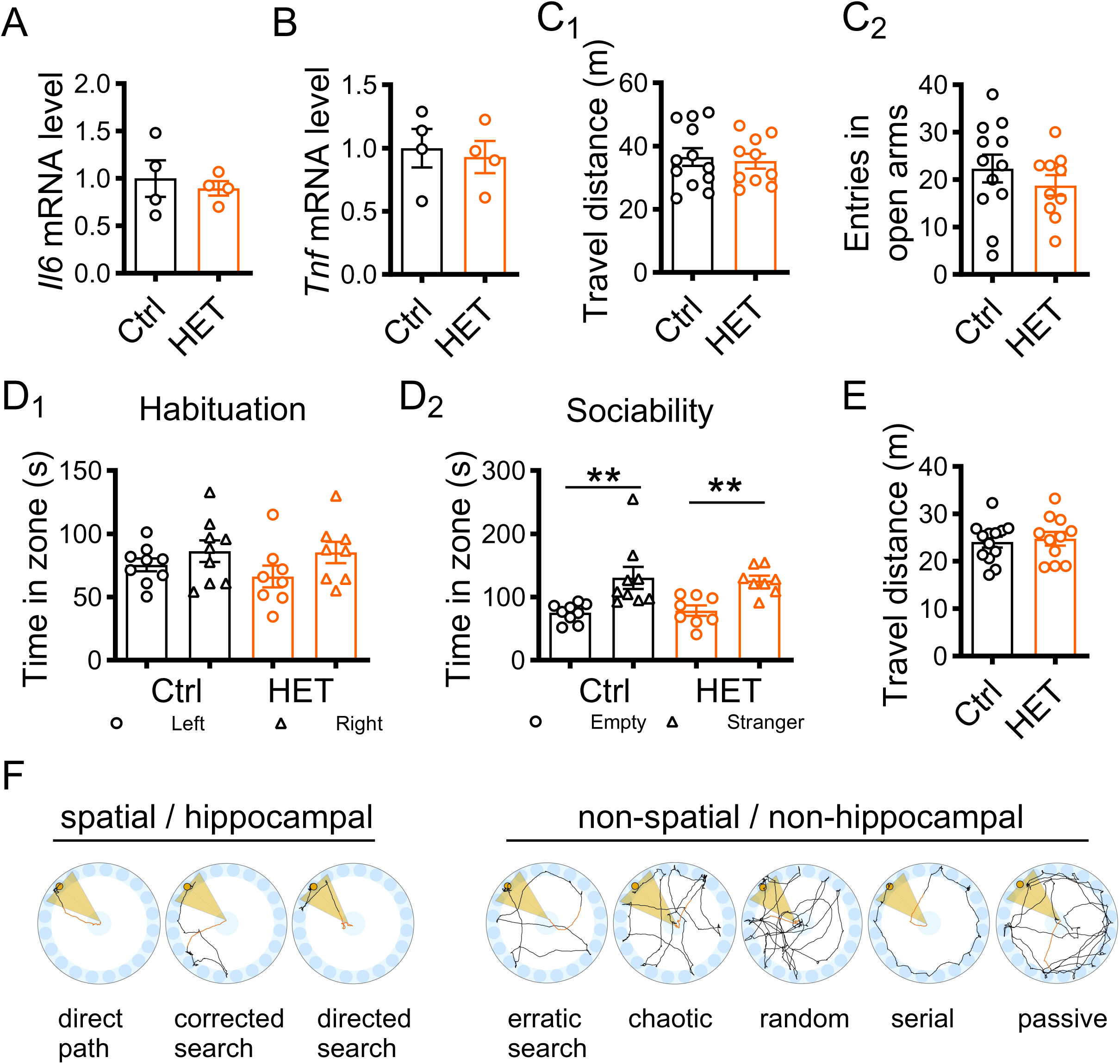
Conditional knockdown of neuronal TAK1 impairs hippocampal function. **A-B**, Hippocampal *Il6* (**A**) and *Tnf* (**B**) mRNA expression in Ctrl and HET mice, normalized to actin and expressed relative to Ctrl (Ctrl: n = 4 mice; HET: n = 4 mice). **C,** Additional elevated plus-maze measurements. (**C_1_**), Total distance travelled. (**C_2_**), Number of entries into the open arms (Ctrl: n = 12 mice; HET: n = 10 mice). **D,** Three-chamber test performance. (**D_1_**), Time spent in the left and right chambers during habituation. (**D_2_**), Time spent investigating the empty enclosure and stranger mouse during the sociability phase (Ctrl: n = 9 mice, P = 0.004; HET: n = 8 mice, P = 0.001; Wilcoxon matched-pairs signed rank test). **E,** Total distance travelled during Y maze (Ctrl: n = 13 mice; HET: n = 11 mice). **F,** Drawings exemplifying color-coded searching trajectories, including non-spatial and spatial strategies. Data are presented as mean ± s.e.m. Statistical tests: Two-sided Mann–Whitney U test was used unless otherwise stated. Exact P values are reported for comparisons indicated in the figure; complete statistical results, including nonsignificant comparisons, are provided in the Key resource table 4. Data summary.

**Supplementary Figure 2.**
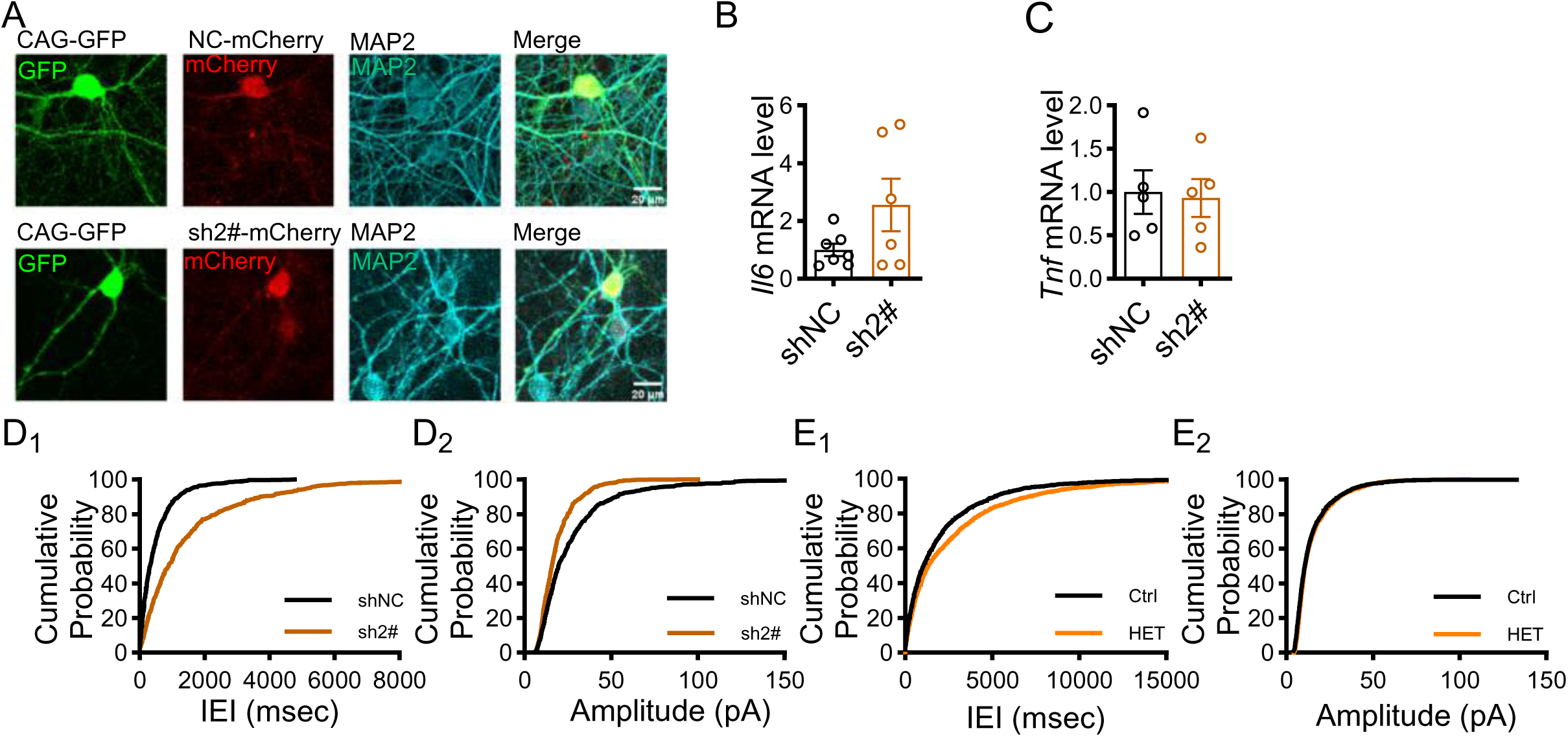
shRNA-mediated neuronal TAK1 knockdown impairs spine structure and synaptic transmission. **A**, Representative images of neurons transfected with CAG-GFP together with negative control mCherry (shNC) or sh2#-mCherry (sh2#) and immunostained for MAP2. Scale bars, 20 μm. **B,** Relative *Il6* mRNA expression in neurons transfected with shNC or sh2# (shNC: n = 7 wells from 2 cultures; sh2#: n = 6 wells from 2 cultures; P = 0.295). **C,** Relative *Tnf* mRNA expression in neurons transfected with shNC or sh2# (shNC: n = 5 wells from 2 cultures; sh2#: n = 5 wells from 2 cultures; P > 0.999). **D,** Cumulative distributions of mEPSC inter-event intervals (**D_1_**) and amplitudes (**D_2_**) in neurons expressing shNC or sh2# (shNC: n = 15 cells from 2 cultures; sh2#: n = 17 cells from 2 cultures; P_IEI < 0.001; P_amplitude < 0.001; Kolmogorov–Smirnov test). **E,** Cumulative distributions of mEPSC inter-event intervals (**E_1_**) and amplitudes (**E_2_**) in CA1 pyramidal neurons from Ctrl and HET mice (Ctrl: n = 19 cells from 6 mice; HET: n = 13 cells from 4 mice; P_IEI < 0.001; P_amplitude < 0.001; Kolmogorov–Smirnov test). Data are presented as mean ± s.e.m. in B and C. Statistical tests: Two-sided Mann– Whitney U test was used unless otherwise stated. Exact P values are reported for comparisons indicated in the figure; complete statistical results, including nonsignificant comparisons, are provided in the Key resource table 4. Data summary.

**Supplementary Figure 3.**
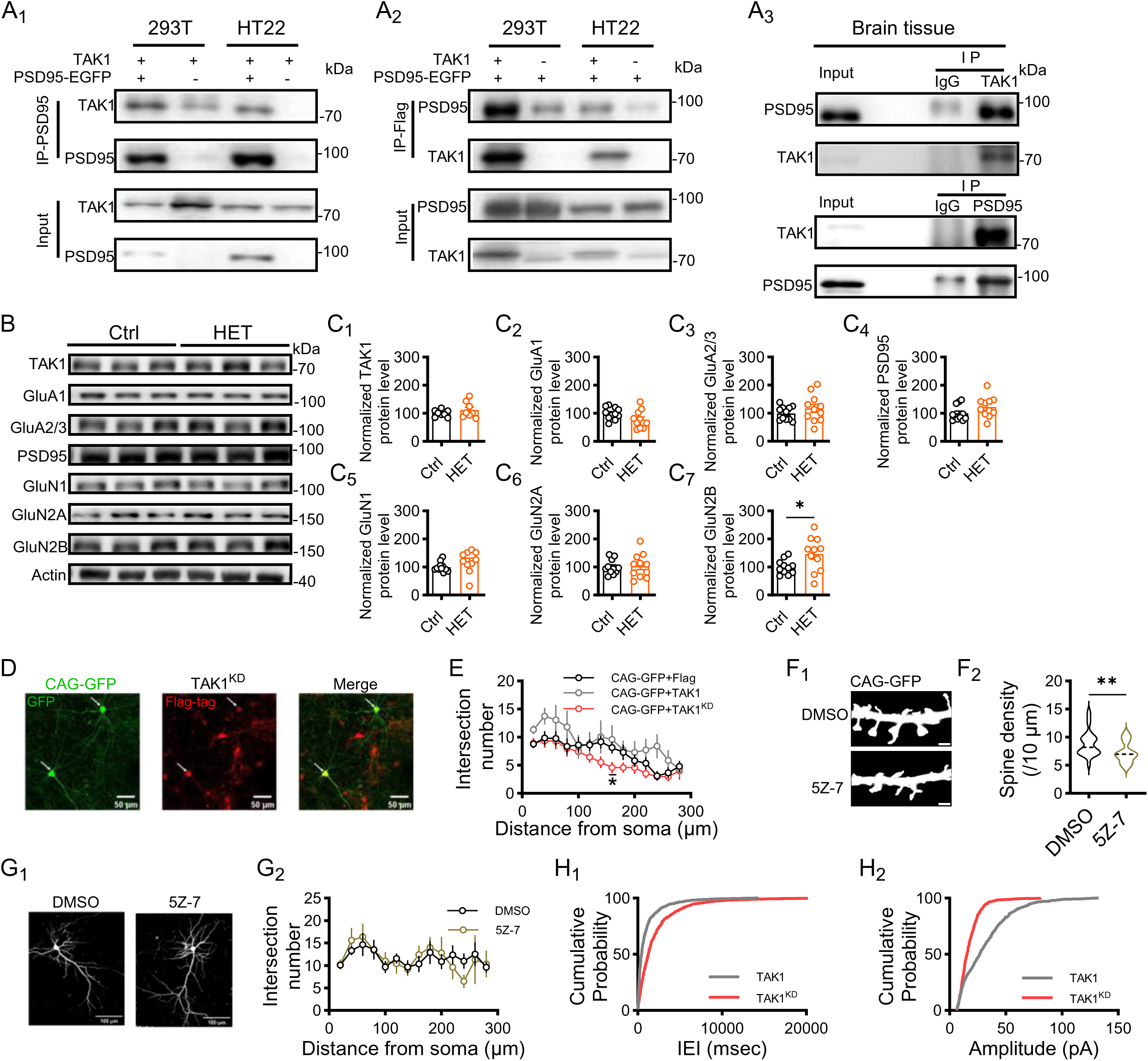
Postsynaptic localization of TAK1 and its kinase activity support excitatory synapse integrity. **A**, Association between TAK1 and PSD95. (**A_1_-A_2_**), Reciprocal co-immunoprecipitation of TAK1 and PSD95-EGFP in HEK293T and HT22 cells. Immunoprecipitation was performed using antibodies against PSD95 (**A_1_**) or Flag (**A_2_**). (**A_3_**), Reciprocal co-immunoprecipitation of endogenous TAK1 and PSD95 from brain tissue. IgG served as the immunoprecipitation control. **B,** Representative immunoblots of TAK1, GluA1, GluA2/3, PSD95, GluN1, GluN2A and GluN2B in hippocampal soluble fractions (S1) from Ctrl and HET mice. Actin served as a loading control. **C,** Quantification of TAK1 (**C_1_**), GluA1 (**C_2_**), GluA2/3 (**C_3_**), PSD95 (**C_4_**), GluN1 (**C_5_**), GluN2A (**C_6_**) and GluN2B (**C_7_**, P = 0.047) in S1, normalized to actin and expressed relative to Ctrl (Ctrl: n = 11 mice; HET: n = 11 mice). **D-E,** Dendritic arborization in neurons co-expressing CAG-GFP with Flag, TAK1 or TAK1^KD^. **D,** Representative images. Arrowheads indicate transfected neurons. Scale bars, 50 μm. **E,** Sholl analysis (n = 12-14 neurons from 2 cultures, Two-way ANOVA with Sidak post hoc multiple-comparisons, F (2,37) = 1.621, P = 0.211; P_CAG-GFP+TAK1 vs. CAG-GFP+ TAK1^KD^ (160 μm) = 0.037). **F,** Dendritic spine density after pharmacological TAK1 inhibition. (**F_1_**), Representative dendritic segments from neurons treated with DMSO or 5Z-7. Scale bars, 2 μm. (**F_2_**), Quantification of spine density (DMSO: n = 24 dendritic segments from 12 neurons and 2 cultures; 5Z-7: n = 15 dendritic segments from 8 neurons and 2 cultures; P = 0.008). **G,** Dendritic arborization after 5Z-7 treatment. (**G_1_**), Representative neurons. Scale bars, 100 μm. (**G_2_**), Sholl analysis (DMSO: n = 9 neurons from 2 cultures; 5Z-7: n = 5 neurons from 2 cultures). **H,** Cumulative distributions of mEPSC inter-event intervals (**H_1_**) and amplitudes (**H_2_**) in neurons expressing TAK1 or TAK1^KD^ (n = 9-10 neurons from 1 culture; P_TAK1 vs. TAK1^KD^ < 0.001; Kolmogorov–Smirnov test). Data are presented as mean ± s.e.m. Statistical tests: Two-sided Mann–Whitney U test was used unless otherwise stated. Exact P values are reported for comparisons indicated in the figure; complete statistical results, including nonsignificant comparisons, are provided in the Key resource table 4. Data summary.

**Supplementary Figure 4.**
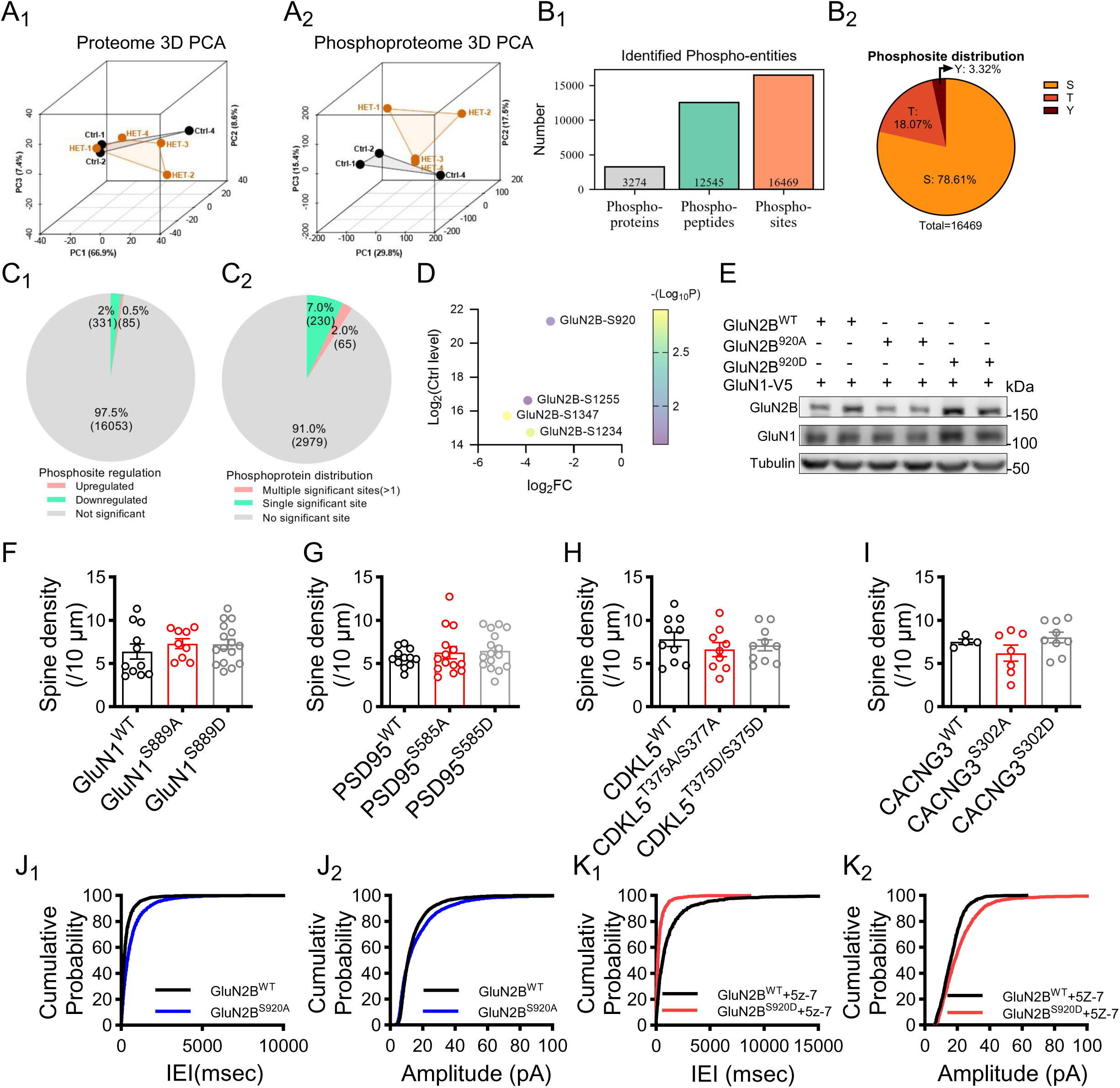
Phosphoproteomic and mutation analysis of functional downstream candidates of neuronal TAK1. **A**, Principal component analysis of the proteome (**A_1_**) and phosphoproteome (**A_2_**). Ctrl-3 was excluded because it exhibited markedly lower phosphopeptide coverage, a substantially higher missing-value rate, the weakest correlation with the other control replicates, and clear separation in PCA. **B,** Summary of the hippocampal PSD phosphoproteomic dataset. (**B_1_**), Numbers of phosphoproteins, phosphopeptides and phosphorylation sites identified. (**B_2_**), Distribution of phosphorylation across serine, threonine and tyrosine residues. **C,** Distribution of regulated phosphosites and phosphoproteins in HET relative to Ctrl mice. (**C_1_**), Proportions of upregulated, downregulated and unchanged phosphosites. (**C_2_**), Proportions of phosphoproteins containing a single regulated phosphosite or multiple regulated phosphosites. Thresholds are defined as |Log_2_FC| > 1 and P value < 0.05. **D,** GluN2B phosphorylation sites plotted by fold change and abundance in Ctrl samples. Circle size represents −log_10_(P). **E,** Co-expression of varied forms of GluN2B (GluN2B^WT^, GluN2B^S920A^, or GluN2B^S920D^) with GluN1-V5 in HEK293T cells. Tubulin served as the loading control. **F–I,** Dendritic spine density in neurons expressing wild-type or phosphosite-mutant forms of GluN1 (**F**), PSD95 (**G**), CDKL5 (**H**) or CACNG3 (**I**). The precise substitutions are indicated beneath each graph (n = 4-16 dendritic segments from 2-8 neurons and 2 cultures). **J,** Cumulative distributions of mEPSC inter-event intervals (**J_1_**) and amplitudes (**J_2_**) in neurons expressing GluN2B^WT^ or GluN2B^S920A^ (n = 12-16 neurons from 2 cultures; P_IEI < 0.001; P_amplitude < 0.001; Kolmogorov–Smirnov test). **K,** Cumulative distributions of mEPSC inter-event intervals (**K_1_**) and amplitudes (**K_2_**) following 5Z-7 treatment in neurons expressing GluN2B^WT^ or GluN2B^S920D^ (n = 7 or 11 neurons from 2 cultures; P_IEI < 0.001; P_amplitude < 0.001; Kolmogorov–Smirnov test). Data are presented as mean ± s.e.m. Statistical tests: Two-sided Mann–Whitney U test was used unless otherwise stated. Exact P values are reported for comparisons indicated in the figure; complete statistical results, including nonsignificant comparisons, are provided in the Key resource table 4. Data summary.

**Supplementary Figure 5.**
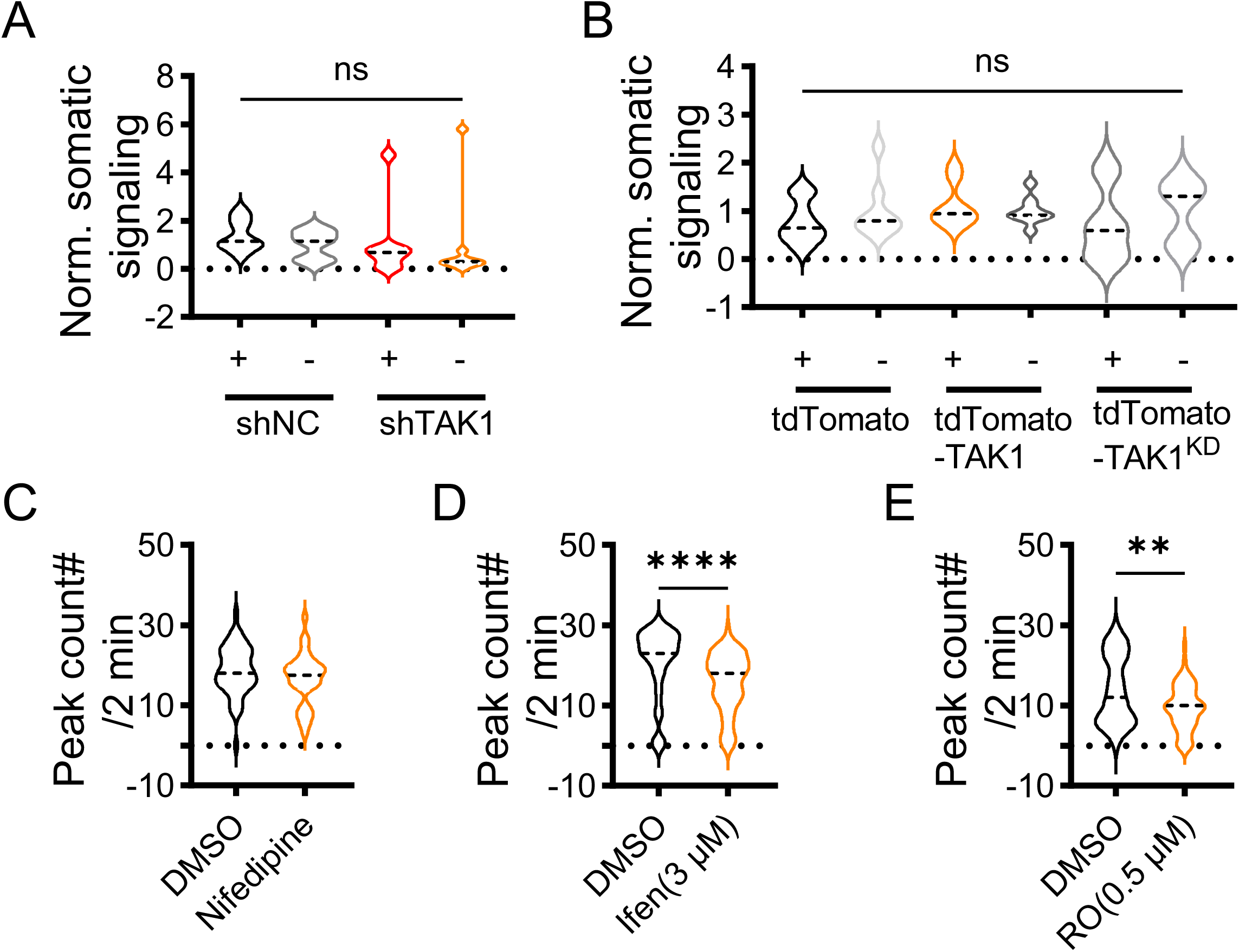
Neuronal TAK1 modulates synaptic but not somatic calcium activity. **A**, Quantification of normalized somatic calcium signals measured with Fluo-8 in shNC or shTAK1-transfected neurons and adjacent untransfected neurons (n = 8 wells from 1 culture). **B,** Quantification of normalized somatic calcium signals in tdTomato neurons, wild-type tdTomato-TAK1 or tdTomato-TAK1^KD^-transfected neurons and adjacent untransfected neurons (n = 8 wells from 1 culture). **C–E,** Number of calcium transients detected over 2 min in neurons treated before and after DMSO or the indicated pharmacological agents: nifedipine (**C**, DMSO, n = 43 synapses from 2 cultures; nifedipine, n = 34 synapses from 2 cultures, P = 0.183), ifenprodil (**D**, n = 59 synapses from 2 cultures, Paired comparisons were analyzed using a two-sided Wilcoxon matched-pairs signed-rank test, P < 0.001), or Ro 25-6981 (**E**, n = 53 synapses from 2 cultures, P = 0.003). Two-sided Wilcoxon matched-pairs signed-rank test. Violin plots show the distribution and central tendency. Statistical tests: Two-sided Mann–Whitney U test was used unless otherwise stated. Exact P values are reported for comparisons indicated in the figure; complete statistical results, including nonsignificant comparisons, are provided in the Key resource table 4. Data summary.

**Supplementary Figure 6.**
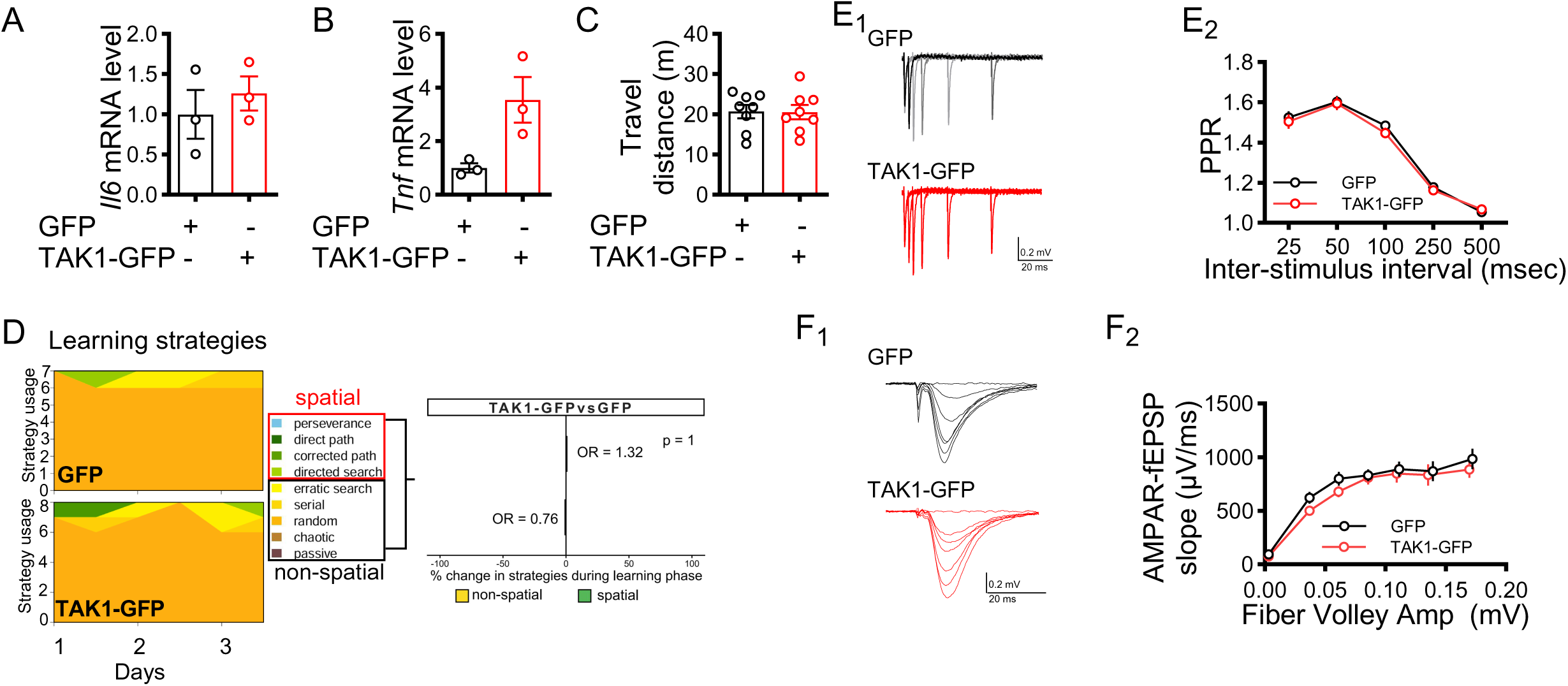
Effects of re-expressing neuronal TAK1 in the hippocampi of aged mice. **A-B**, Relative hippocampal *Il6* (**A**) and *Tnf* (**B**) mRNA expression in aged mice expressing AAV2/9-hSyn-EGFP-WPRE-pA (GFP) or AAV2/9-hSyn-3xFlag-eGFP-mMap3k7-WPRE-pA (TAK1-GFP) (GFP: n = 3 mice; TAK1-GFP: n = 3 mice). **C,** Total distance travelled by aged mice expressing GFP or TAK1-GFP during the Y maze (GFP: n = 8 mice; TAK1-GFP: n = 8 mice). **D,** Search-strategy analysis during Barnes maze acquisition. The left panels show the daily proportions of spatial and non-spatial strategies; the right panel shows the odds ratio for a change in strategy use during acquisition (GFP: n = 7 mice; TAK1-GFP: n = 8 mice). **E,** Paired-pulse ratio at Schaffer collateral–CA1 synapses in aged mice expressing GFP or TAK1-GFP. (**E_1_**), Representative traces. (**E_2_**), Paired-pulse ratio at the indicated interstimulus intervals (GFP: n = 25 slices from 4 mice; TAK1-GFP: n = 32 slices from 4 mice). **F,** Basal excitatory synaptic transmission at Schaffer collateral–CA1 synapses. (**F_1_**), Representative fEPSP waveforms recorded at increasing stimulus intensities. (**F_2_**), Input–output relationship between fiber-volley amplitude and AMPAR-mediated fEPSP slope (GFP: n = 25 slices from 4 mice; TAK1-GFP: n = 32 slices from 4 mice). Data are presented as mean ± s.e.m. Statistical tests: Two-sided Mann–Whitney U test was used unless otherwise stated. Exact P values are reported for comparisons indicated in the figure; complete statistical results, including nonsignificant comparisons, are provided in the Key resource table 4. Data summary.

