## Supplementary Tables for "Postsynaptic TAK1-GluN2B axis supports hippocampal synaptic integrity and cognitive function during aging"

**Key resource table 1. Plasmids, lentiviral vectors and AAV constructs**

| Full name (abbreviation) | Type | Application |
|---|---|---|
| CAG-gag-GFP(CAG-GFP) | plasmid | Neuronal morphology and spine density in vitro |
| U6-shNC/shTAK1-mCherry | plasmid | Neuronal morphology and spine density in vitro |
| CMV-Flag/TAK1$^{KD}$-317-Flag | plasmid | Neuronal morphology and spine density in vitro |
| CMV-TAK1-317-Flag | plasmid | Neuronal morphology, spine density and CO-IP in vitro |
| CMV-tdTomato/tdTomato-TAK1/tdTomato-TAK1$^{KD}$ | plasmid | Synaptic calcium imaging |
| CMV-GluN2B$^{WT}$-V5 | plasmid | Spine density in vitro |
| CMV-GluN2B$^{doubleA}$-V5 | plasmid | Spine density in vitro |
| CMV-GluN2B$^{S920A}$-V5 | plasmid | Spine density in vitro |
| CMV-GluN2B$^{S1255A}$-V5 | plasmid | Spine density in vitro |
| CMV-GluN2B$^{S920D}$-V5 | plasmid | Spine density in vitro |
| CMV-GluN2B$^{S1255D}$-V5 | plasmid | Spine density in vitro |
| CMV-GluN1$^{WT}$-V5 | plasmid | Spine density in vitro |
| CMV-GluN1$^{S889A}$-V5 | plasmid | Spine density in vitro |
| CMV-GluN1$^{S889D}$-V5 | plasmid | Spine density in vitro |
| CMV-PSD95$^{WT}$-V5 | plasmid | Spine density in vitro |
| CMV-PSD95$^{S585A}$-V5 | plasmid | Spine density in vitro |
| CMV-PSD95$^{S585D}$-V5 | plasmid | Spine density in vitro |
| CMV-Cdkl5$^{WT}$-V5 | plasmid | Spine density in vitro |
| CMV-Cdkl5$^{T375A/S377A}$-V5 | plasmid | Spine density in vitro |
| CMV-Cdkl5$^{T375D/S377D}$-V5 | plasmid | Spine density in vitro |
| CMV-Cacng3$^{WT}$-V5 | plasmid | Spine density in vitro |

| | | |
|---|---|---|
| CMV-Cacng3$^{S302A}$-V5 | plasmid | Spine density in vitro |
| CMV-Cacng3$^{S302D}$-V5 | plasmid | Spine density in vitro |
| fast variant of the mNeonGreen-based calcium indicator (NEMOf) | plasmid | Synaptic calcium imaging |
| CMV-GluN2B$^{WT}$-tdTomato | plasmid | Synaptic calcium imaging |
| CMV-GluN2B$^{doubleA}$-tdTomato | plasmid | Synaptic calcium imaging |
| CMV-GluN2B$^{S920A}$-tdTomato | plasmid | Synaptic calcium imaging |
| CMV-GluN2B$^{S1255A}$-tdTomato | plasmid | Synaptic calcium imaging |
| CMV-GluN2B$^{S920D}$-tdTomato | plasmid | Synaptic calcium imaging |
| CMV-GluN2B$^{S1255D}$-tdTomato | plasmid | Synaptic calcium imaging |
| CMV-PSD95-EGFP(PSD95-EGFP) | plasmid | CO-IP in vitro |
| hSyn-GFP-Tet-off (Lenti-hSyn-GFP) | lentivirus | Spine density in vivo and whole cell recording |
| U6-shNC/shTAK1-mCherry | lentivirus | Whole cell recording and somatic calcium imaging |
| CMV-Flag/TAK1-317-Flag/TAK1$^{KD}$-317-Flag | lentivirus | Whole cell recording |
| CMV-GluN2B$^{WT}$-V5 | lentivirus | Whole cell recording |
| CMV-GluN2B$^{S920A}$-V5 | lentivirus | Whole cell recording |
| CMV-GluN2B$^{S920D}$-V5 | lentivirus | Whole cell recording |
| CMV-tdTomato/tdTomato-TAK1/tdTomato-TAK1$^{KD}$ | lentivirus | somatic calcium imaging |
| AAV2/9-hSyn-3xFlag-eGFP-mMap3k7-WPRE-pA (TAK1-GFP) | AAV | IF in vitro and injected in aging mice |
| AAV2/9-hSyn-EGFP-WPRE-pA (GFP) | AAV | Injected in aging mice |

**Key resource table 2. Main Reagents**

| Name | Source | Catalog NO. | Application | Concentration used |
|---|---|---|---|---|
| 5-fluoro-2'-deoxyuridine | Sigma | Cat# F0503 | Primary rat neuron culture | 8 μM |
| Uridine | Sigma | Cat# U3003 | Primary rat neuron culture | 20 μM |
| Lipo3000 | Thermo Fisher | Cat# L3000015 | Transfection of plasmids into primary neurons | 1μl or 1.2μl per well |
| Neurobasal™ Medium | Gibco | Cat# 21103049 | Primary neuron culture | |
| B-27™ Supplement (50X) | Gibco | Cat# 17504001 | Primary neuron culture | 2 % |
| GlutaMAX | Gibco | Cat# 35050061 | Primary neuron culture | 0.25 % |
| 5Z-7-Oxozeaenol | Sigma | Cat# 09890 | Neuronal morphology, spine density and synaptic calcium imaging in vitro | 300 nM |
| Fluo-8 AM | Maokang Biotechnology | Cat# MX4505 | Somatic calcium imaging | 4 μM |
| Nifedipine | Sigma | Cat# 481981 | Synaptic calcium imaging in vitro | 8 μM |
| Ro 25-6981 | Abcam | Cat# ab120290 | Synaptic calcium imaging in vitro | 0.5 μM |
| Ifenprodil | Sigma | Cat# 23210-58-4 | Synaptic calcium imaging in vitro | 3 μM |
| Tetrodotoxin (TTX) | Sigma-Aldrich | Cat# 4368-28-9 | Whole cell recording | 1 μM |

| | | | | |
|---|---|---|---|---|
| D-APV | Tocris | Cat# 0105 | Whole cell recording | 100 μM |
| SR-95531 (gabazine) | Tocris | Cat# 1262 | Whole cell recording | 10 μM |
| Flag-Affinity Gel | Bimake | Cat# B23102 | CO-IP in cell lines | 10 ul/tube |
| Protein G Magnetic Beads | Thermo Fisher | Cat# 88847 | CO-IP in mouse brain tissue | 10 ul/tube |

**Key resource table 3. Antibodies used for western blotting and immunostaining**

| Antibodies | Source | Catalog NO. | Application | Dilution |
|---|---|---|---|---|
| DAPI | Thermo Fisher | Cat# D1306 | IF | 1:500 |
| MAP2 | Abcam | Cat# Ab5392 | IF | 1:2000 |
| TAK1 | Abcam | Cat# Ab109526 | WB | 1:1000 |
| | | | IF | 1:200 |
| | | | CO-IP | 3 ug/mg protein |
| V5-tag | Invitrogen | Cat# R960-25 | WB/IF | 1:1000 |
| mCherry | Proteintech | Cat# 26765-1-AP | IF | 1:800 |
| p-Thr184/187 TAK1 | Cell Signaling Technology | Cat# 4508 | WB | 1:1000 |
| Normal mouse IgG | Santa Cruz | Cat# sc-2025 | CO-IP | 2.75 ug/mg protein |
| Normal rabbit IgG | Cell Signaling Technology | Cat# 2729s | CO-IP | 3 ug/mg protein |
| GluA1 | Millipore | Cat# MAB2263 | WB | 1:1000 |
| GluA2/3 | Millipore | Cat# AB1506 | WB | 1:1000 |
| GluN1, CT | Millipore | Cat# 05-432 | WB | 1:1000 |

| | | | | |
|---|---|---|---|---|
| GluN2A | Millipore | Cat# 07-632 | WB | 1:1000 |
| GluN2B, clone BWJHL | Millipore | Cat# 05-920 | WB | 1:1000 |
| PSD-95 (K28/43) | Neuromab | Cat# 75-028 | CO-IP | 2.75 ug/mg protein |
| | | | WB | 1:5000 |
| β-actin | Santa Cruz | Cat# sc-47778 | WB | 1:4000 |
| α-Tubulin pAb | MBL | Cat# PM054 | WB | 1:40000 |
| GFP | Abcam | Cat# 13970 | IF | 1:1000 |
| DYKDDDDK Tag (Flag-tag) | Thermo Fisher | Cat# MA5-50638 | IF and WB | 1:1000 |
| Goat Anti-mouse IgG H&L (Alexa Fluor® 488) | Abcam | Cat# ab 150113 | IF | 1:1000 |
| Goat Anti-chicken IgG H&L (Alexa Fluor® 488) | Thermo Fisher | Cat# A10039 | IF | 1:1000 |
| Goat Anti-Rabbit IgG H&L (Alexa Fluor® 568) | Abcam | Cat# ab175471 | IF | 1:1000 |
| Goat Anti-chicken IgG H&L (Alexa Fluor® 647) | Thermo Fisher | Cat# A21449 | IF | 1:1000 |
| Peroxidase-Conjugated Goat Anti-Mouse IgG(H+L) | Yeasen Biotech | Cat# 33201ES60 | WB | 1:10000 |
| Peroxidase-Conjugated Goat Anti-Rabbit IgG(H+L) | Yeasen Biotech | Cat# 33101ES60 | WB | 1:10000 |

**Key resource table 4. Data summary**

| Figure | Group | Mean | N | SEM | P value |
|---|---|---|---|---|---|
| 1B | Ctrl | 25 | 14 | 0.78 | 0.768 |
| | HET | 24.5 | 11 | 0.88 | |
| 1C | Ctrl | 1 | 10 | 0.07 | 0.003 |
| | HET | 0.71 | 6 | 0.32 | |
| $1D_2$ | Ctrl | 44.19 | 13 | 2.96 | 0.232 |
| | HET | 40.5 | 10 | 2.05 | |
| $1D_3$ | Ctrl | 56.17 | 12 | 2.65 | 0.786 |
| | HET | 55.23 | 11 | 5.16 | |
| $1E_2$ | Ctrl | 9.99 | 12 | 1.76 | 0.221 |
| | HET | 7.22 | 10 | 1.09 | |
| $1F_2$ | Ctrl-familiar mouse | 76.33 | 9 | 8.53 | Wilcoxon matched-pairs signed rank test P=0.004 |
| | Ctrl-new mouse | 130.68 | 9 | 5.19 | |
| | HET-familiar mouse | 66.76 | 8 | 6.52 | Wilcoxon matched-pairs signed rank test P=0.016 |
| | HET-new mouse | 111.59 | 8 | 12.23 | |
| $1G_2$ | Ctrl | 62.34 | 12 | 2.52 | 0.046 |
| | HET | 52.38 | 11 | 2.49 | |
| $1H_3$ | Ctrl-D1 | 97.25 | 12 | 5.68 | unpaired t-test，P_Ctrl-D1 vs. HET-D1 = 0.562, P_Ctrl-D2 vs. HET-D2 = 0.847, P_Ctrl-D3 vs. HET-D3 = 0.759,P_Ctrl-D4 vs. HET-D4 = 0.032,P_Ctrl-D5 vs. HET-D5 = 0.302, P_Ctrl-D6 vs. HET-D6 = 0.977 |
| | HET-D1 | 111.36 | 9 | 4.9 | |
| | Ctrl-D2 | 76.41 | 12 | 3.72 | |
| | HET-D2 | 80.41 | 9 | 6.72 | |
| | Ctrl-D3 | 41.57 | 11 | 4.12 | |
| | HET-D3 | 46.28 | 9 | 5.98 | |
| | Ctrl-D4 | 24.65 | 11 | 3.74 | |
| | HET-D4 | 52.75 | 8 | 4.63 | |
| | Ctrl-D5 | 34.28 | 12 | 4.47 | |
| | HET-D5 | 22.95 | 9 | 4.45 | |
| | Ctrl-D6 | 39.18 | 12 | 4.7 | |
| | HET-D6 | 39.55 | 9 | 4.4 | |
| $1H_4$ | Ctrl | 53.91 | 12 | 3.22 | 0.041 |
| | HET | 43.53 | 9 | 2.84 | |
| $1J_2$ | Ctrl-25ms | 1.58 | 31 | 0.03 | unpaired t-test with Welch correction, P_Ctrl-25 ms vs. HET-25 ms = 0.019, P_Ctrl-50 ms vs. HET-50 ms = 0.044, P_Ctrl-100 ms vs. HET-100 ms = |
| | HET-25ms | 1.48 | 31 | 0.03 | |
| | Ctrl-50ms | 1.74 | 31 | 0.04 | |
| | HET-50ms | 1.63 | 31 | 0.03 | |
| | Ctrl-100ms | 1.52 | 31 | 0.02 | |
| | HET-100ms | 1.46 | 31 | 0.02 | |

| | | | | | |
|---|---|---|---|---|---|
| | Ctrl-250ms | 1.24 | 31 | 0.02 | 0.05, P_Ctrl-250 ms vs. HET-250 ms = 0.08, P_Ctrl-500 ms vs. HET-500 ms = 0.71, |
| | HET-250ms | 1.19 | 31 | 0.02 | |
| | Ctrl-500ms | 1.07 | 31 | 0.01 | |
| | HET-500ms | 1.06 | 31 | 0.01 | |
| $1K_2$ | Ctrl-0.01mV | 394.64 | 33 | 34.98 | unpaired t-test with Welch correction, P_Ctrl-0.01 mV vs. HET-0.01 mV = 0.637, P_Ctrl-0.04 mV vs. HET-0.04 mV = 0.378, P_Ctrl-0.06 mV vs. HET-0.06 mV = 0.367, P_Ctrl-0.09 mV vs. HET-0.09 mV = 0.09, P_Ctrl-0.11 mV vs. HET-0.11 mV = 0.325, P_Ctrl-0.13 mV vs. HET-0.13 mV = 0.539, P_Ctrl-0.16 mV vs. HET-0.16 mV = 0.715 |
| | HET-0.01mV | 370.99 | 30 | 35.48 | |
| | Ctrl-0.04mV | 563.34 | 32 | 44.01 | |
| | HET-0.04mV | 515.76 | 31 | 30.56 | |
| | Ctrl-0.06mV | 690.47 | 36 | 42.47 | |
| | HET-0.06mV | 741.65 | 32 | 37.09 | |
| | Ctrl-0.09mV | 789.89 | 35 | 48.84 | |
| | HET-0.09mV | 899.41 | 32 | 40.71 | |
| | Ctrl-0.11mV | 932.33 | 35 | 58.74 | |
| | HET-0.11mV | 1010.75 | 32 | 52.93 | |
| | Ctrl-0.13mV | 1025.48 | 28 | 72.02 | |
| | HET-0.13mV | 1081.34 | 30 | 54.46 | |
| | Ctrl-0.16mV | 1073.88 | 19 | 96.6 | |
| | HET-0.16mV | 1118.48 | 19 | 73.31 | |
| $1L_2$ | Ctrl | 134.32 | 22 | 7.67 | 0.001 |
| | HET | 96.68 | 21 | 6.90 | |
| $1M_2$ | Ctrl | 79.17 | 9 | 4.72 | 0.554 |
| | HET | 76.94 | 12 | 3.98 | |
| 2B | shNC | 100 | 3 | 0 | 0.03 |
| | Sh1# | 53.7 | 3 | 8.41 | |
| | shNC | 100 | 6 | 0 | < 0.001 |
| | Sh2# | 41.58 | 4 | 6.66 | |
| | shNC | 100 | 3 | 0 | 0.377 |
| | Sh3# | 88.88 | 4 | 10.76 | |
| $2C_2$ | ShNC-20μm | 10.24 | 10 | 1.49 | 0.436 |
| | Sh2#-20μm | 11.51 | 10 | 1.14 | |
| | ShNC-40μm | 15.5 | 10 | 3.45 | 0.393 |
| | Sh2#-40μm | 15.87 | 10 | 1.72 | |
| | ShNC-60μm | 14.63 | 10 | 2.38 | 0.218 |
| | Sh2#-60μm | 16.71 | 10 | 1.82 | |
| | ShNC-80μm | 13.08 | 10 | 1.9 | 0.353 |
| | Sh2#-80μm | 14.95 | 10 | 1.84 | |
| | ShNC-100μm | 12.21 | 10 | 1.77 | 0.796 |
| | Sh2#-100μm | 11.54 | 10 | 1.92 | |
| | ShNC-120μm | 11.77 | 10 | 2.2 | 0.28 |
| | Sh2#-120μm | 8.55 | 10 | 1.42 | |

| | | | | | |
|---|---|---|---|---|---|
| | ShNC-140μm | 9.64 | 10 | 2.31 | 0.739 |
| | Sh2#-140μm | 7.2 | 10 | 1.01 | |
| | ShNC-160μm | 7.71 | 10 | 2.12 | 0.853 |
| | Sh2#-160μm | 6.54 | 10 | 1.3 | |
| | ShNC-180μm | 7.13 | 8 | 2.55 | 0.965 |
| | Sh2#-180μm | 6.06 | 10 | 1.16 | |
| | ShNC-200μm | 7.51 | 6 | 2.12 | 0.145 |
| | Sh2#-200μm | 3.68 | 9 | 0.9 | |
| | ShNC-220μm | 4.12 | 6 | 1.29 | 0.662 |
| | Sh2#-220μm | 3.22 | 8 | 0.7 | |
| | ShNC-240μm | 3.25 | 5 | 0.99 | > 0.999 |
| | Sh2#-240μm | 3.33 | 5 | 0.63 | |
| | ShNC-260μm | 1.7 | 3 | 0.35 | 0.629 |
| | Sh2#-260μm | 2.58 | 4 | 0.57 | |
| $2D_2$ | ShNC | 9.87 | 33 | 0.36 | < 0.001 |
| | Sh2# | 6.69 | 28 | 0.42 | |
| $2D_3$ | ShNC | 2.13 | 674 | 0.06 | K-S test, P > 0.999 |
| | Sh2# | 1.71 | 392 | 0.07 | |
| $2E_2$ | ShNC | 2.17 | 15 | 0.25 | < 0.001 |
| | Sh2# | 0.85 | 17 | 0.11 | |
| $2E_3$ | ShNC | 28.85 | 15 | 3.75 | 0.076 |
| | Sh2# | 19.95 | 17 | 1.8 | |
| $2F_3$ | Ctrl | 18.31 | 31 | 0.8 | < 0.001 |
| | HET | 12.8 | 40 | 0.54 | |
| $2F_4$ | Ctrl | 2.15 | 780 | 0.04 | K-S test, P > 0.999 |
| | HET | 1.84 | 679 | 0.04 | |
| $2G_2$ | Ctrl | 0.57 | 19 | 0.04 | 0.002 |
| | HET | 0.38 | 13 | 0.03 | |
| $2G_3$ | Ctrl | 14.52 | 19 | 0.8 | 0.158 |
| | HET | 15.72 | 13 | 0.71 | |
| $3D_1$ | Ctrl | 97.96 | 6 | 6.67 | 0.004 |
| | HET | 67.27 | 6 | 5.86 | |
| $3D_2$ | Ctrl | 100.4 | 6 | 2.79 | 0.725 |
| | HET | 101.6 | 6 | 7.57 | |
| $3D_3$ | Ctrl | 99.83 | 6 | 3.53 | 0.221 |

| | | | | | |
|---|---|---|---|---|---|
| | HET | 113.17 | 6 | 8.23 | |
| $3D_4$ | Ctrl | 100.17 | 6 | 5.08 | 0.333 |
| | HET | 92.5 | 6 | 3.56 | |
| $3D_5$ | Ctrl | 100 | 6 | 4.13 | 0.513 |
| | HET | 105 | 6 | 4.64 | |
| $3D_6$ | Ctrl | 100 | 6 | 7.55 | 0.558 |
| | HET | 100.83 | 6 | 5.31 | |
| $3G_2$ | Flag | 9.01 | 24 | 0.49 | ANOVA $P < 0.001$, $F(2, 64) = 10.17$, P_Flag vs Flag-TAK1 = 0.959 |
| | TAK1 | 9.22 | 21 | 0.55 | |
| | $TAK1^{KD}$ | 6.22 | 22 | 0.53 | |
| $3G_3$ | Flag | 2.76 | 393 | 0.1 | K-S test P_Flag vs Flag-TAK1 = 0.251 |
| | TAK1 | 1.63 | 237 | 0.08 | |
| | $TAK1^{KD}$ | 2.55 | 360 | 0.1 | |
| $3H_2$ | TAK1 | 2.24 | 10 | 0.68 | 0.044 |
| | $TAK1^{KD}$ | 0.69 | 9 | 0.14 | |
| $3H_3$ | TAK1 | 29.24 | 10 | 4.12 | 0.113 |
| | $TAK1^{KD}$ | 18.49 | 9 | 2.04 | |
| 4E | $GluN2B^{WT}$ | 8.06 | 16 | 0.53 | ANOVA_P < 0.001, $F(3, 50) = 16.53$, P_$GluN2B^{WT}$ vs. $GluN2B^{S1255A}$ = 0.042, P_$GluN2B^{S920A}$ vs. $GluN2B^{doubleA}$ = 0.910 |
| | $GluN2B^{S920A}$ | 5.44 | 12 | 0.33 | |
| | $GluN2B^{S1255A}$ | 10.04 | 9 | 0.76 | |
| | $GluN2B^{doubleA}$ | 5.88 | 17 | 0.32 | |
| $4F_2$ | $GluN2B^{WT}$ | 3.72 | 16 | 0.43 | 0.004 |
| | $GluN2B^{S920A}$ | 1.82 | 12 | 0.26 | |
| $4F_3$ | $GluN2B^{WT}$ | 13.65 | 16 | 1.14 | 0.397 |
| | $GluN2B^{S920A}$ | 17.44 | 12 | 2.51 | |
| 4G | ShTAK1+ $GluN2B^{WT}$ | 5.69 | 19 | 0.39 | ANOVA_P < 0.001, $F(2, 39) = 9.414$, P_shTAK1+$GluN2B^{WT}$ vs. shTAK1+$GluN2B^{S1255D}$ = 0.082, P_shTAK1+$GluN2B^{S920D}$ vs. shTAK1+$GluN2B^{S1255D}$ = 0.104 |
| | ShTAK1+ $GluN2B^{S920D}$ | 8.21 | 10 | 0.4 | |
| | ShTAK1+ $GluN2B^{S1255D}$ | 6.89 | 13 | 0.38 | |
| $4H_2$ | 5Z-7+ $GluN2B^{WT}$ | 0.85 | 7 | 0.17 | 0.003 |
| | 5Z-7+ $GluN2B^{S920D}$ | 3.56 | 11 | 0.5 | |
| $4H_3$ | 5Z-7+ $GluN2B^{WT}$ | 15.6 | 7 | 1.39 | 0.179 |

| | | | | | |
|---|---|---|---|---|---|
| | 5Z-7+ GluN2B$^{S920D}$ | 21.39 | 11 | 2.42 | |
| 5B | ShNC | 15.26 | 110 | 0.6 | < 0.001 |
| | ShTAK1 | 9.32 | 98 | 0.69 | |
| 5C | tdTomato | 13.91 | 117 | 0.66 | ANOVA_P < 0.001, F (2, 346) = 28.13, P_dTomato vs. TAK1-dTomato = 0.981 |
| | tdTomato-TAK1$^{KD}$ | 14.12 | 100 | 0.75 | |
| | tdTomato-TAK1 | 7.3 | 132 | 0.81 | |
| 5D | DMSO | 16.7 | 125 | 0.63 | 0.027 |
| | 5Z-7 | 14.2 | 157 | 0.75 | |
| 5E | GluN2B$^{WT}$ | 15.13 | 125 | 0.4 | ANOVA_P < 0.001, F (3, 446) = 12.89, P_GluN2B vs. S1255A = 0.085, Double A vs. S920A = 0.888, |
| | GluN2B$^{doubleA}$ | 12.63 | 120 | 0.67 | |
| | GluN2B$^{S920A}$ | 11.9 | 87 | 0.98 | |
| | GluN2B$^{S1255A}$ | 17.31 | 118 | 0.7 | |
| 5F | DMSO | 14.81 | 37 | 1.16 | ANOVA_P < 0.001, F (3, 154) = 9.038, P_S920D vs. DMSO = 0.221, P_S1255D vs. DMSO = 0.244, P_GluN2B vs. S1255D = 0.952 |
| | 5Z-7+GluN2B$^{WT}$ | 11.25 | 51 | 0.82 | |
| | 5Z-7+GluN2B$^{S920D}$ | 17.75 | 40 | 1.19 | |
| | 5Z-7+GluN2B$^{S1255D}$ | 11.73 | 30 | 1.26 | |
| $6A_2$ | Young | 100 | 7 | 5.41 | 0.639 |
| | Aged | 95.29 | 5 | 6.4 | |
| $6B_2$ | Young | 100 | 6 | 5.02 | 0.004 |
| | Aged | 79.72 | 5 | 1.3 | |
| $6D_2$ | GFP | 42.36 | 8 | 3.52 | 0.038 |
| | TAK1-GFP | 55.1 | 8 | 3.88 | |
| $6E_2$ | D1-GFP | 147.64 | 7 | 2.62 | unpaired t-test with Welch correction, P_D1-GFP vs. D1-TAK1-GFP = 0.525, P_D2-GFP vs. D2-TAK1-GFP = 0.701, P_D3-GFP vs. D3-TAK1-GFP = 0.889, P_D4-GFP vs. D4-TAK1-GFP = 0.427, P_D5-GFP vs. D5-TAK1- |
| | D1-TAK1-GFP | 133.13 | 8 | 4.56 | |
| | D2-GFP | 142.93 | 7 | 3.94 | |
| | D2-TAK1-GFP | 133.88 | 8 | 3.58 | |
| | D3-GFP | 133.57 | 7 | 5.06 | |
| | D3-TAK1-GFP | 129.69 | 8 | 3.98 | |
| | D4-GFP | 123.5 | 7 | 3.77 | |

| | | | | | |
|---|---|---|---|---|---|
| | D4-TAK1-GFP | 103.31 | 8 | 5.25 | GFP = 0.032, P_D6-GFP vs. D6-TAK1-GFP = 0.037 |
| | D5-GFP | 127.43 | 7 | 6 | |
| | D5-TAK1-GFP | 56.43 | 7 | 2.95 | |
| | D6-GFP | 118.93 | 7 | 4.58 | |
| | D6-TAK1-GFP | 65.75 | 8 | 4.07 | |
| $6E_3$ | GFP | 100 | 7 | 9.15 | 0.022 |
| | TAK1-GFP | 142.93 | 6 | 10.57 | |
| $6E_4$ | GFP | 160.43 | 7 | 12.77 | < 0.001 |
| | TAK1-GFP | 39.29 | 7 | 8.67 | |
| $6F_2$ | GFP | 13.05 | 17 | 0.73 | 0.03 |
| | TAK1-GFP | 15.78 | 15 | 0.7 | |
| $6G_2$ | GFP | 120.87 | 10 | 8.19 | Welch's test P = 0.032 |
| | TAK1-GFP | 149.72 | 12 | 9.43 | |
| S1A | Ctrl | 1 | 4 | 0.19 | 0.886 |
| | HET | 0.9 | 4 | 0.08 | |
| S1B | Ctrl | 1 | 4 | 0.15 | 0.686 |
| | HET | 0.93 | 4 | 0.13 | |
| $S1C_1$ | Ctrl | 36.53 | 12 | 2.83 | 0.821 |
| | HET | 35.23 | 10 | 2.36 | |
| $S1C_2$ | Ctrl | 22.33 | 12 | 2.92 | 0.275 |
| | HET | 18.8 | 10 | 2.15 | |
| $S1D_1$ | Ctrl-left | 75.5 | 9 | 5.03 | Wilcoxon matched-pairs signed rank test, P = 0.098 |
| | Ctrl-right | 86.29 | 9 | 8.53 | |
| | HET-left | 66.26 | 8 | 8.6 | Wilcoxon matched-pairs signed rank test,P = 0.134 |
| | HET-right | 85.2 | 8 | 8.52 | |
| $S1D_2$ | Ctrl-empty | 75.17 | 9 | 4.86 | 0.004 |
| | Ctrl-stranger1 | 130.47 | 9 | 17.39 | |
| | HET-empty | 78.38 | 8 | 8.25 | 0.008 |
| | HET-stranger1 | 126.06 | 8 | 7.43 | |
| S1E | Ctrl | 24.1 | 13 | 1.14 | 0.776 |
| | HET | 24.78 | 11 | 1.42 | |
| S2B | Ctrl | 1 | 7 | 0.22 | 0.295 |
| | HET | 2.56 | 6 | 0.91 | |
| S2C | Ctrl | 1 | 5 | 0.25 | >0.999 |
| | HET | 0.93 | 5 | 0.22 | |
| $S2D_1$ | shNC | 515.52 | 750 | 22.31 | K-S test P < 0.001 |
| | Sh2# | 1551.79 | 815 | 79.11 | |

| | | | | | |
|---|---|---|---|---|---|
| $S2D_2$ | shNC | 27.78 | 750 | 0.89 | K-S test P < 0.001 |
| | Sh2 # | 18.72 | 815 | 0.37 | |
| $S2E_1$ | Ctrl | 1954.423 | 3699 | 120.5828 | K-S test P < 0.001 |
| | HET | 2816.234 | 1975 | 92.31772 | |
| $S2E_2$ | Ctrl | 14.53522 | 3699 | 0.208616 | K-S test P < 0.001 |
| | HET | 15.95982 | 1975 | 0.27924 | |
| $S3C_1$ | Ctrl | 100 | 8 | 4.38 | 0.645 |
| | HET | 111.95 | 8 | 10.05 | |
| $S3C_2$ | Ctrl | 100 | 11 | 6.43 | 0.076 |
| | HET | 77.57 | 11 | 9.44 | |
| $S3C_3$ | Ctrl | 100 | 11 | 7.37 | 0.478 |
| | HET | 117.26 | 11 | 14.61 | |
| $S3C_4$ | Ctrl | 100 | 11 | 8.41 | 0.076 |
| | HET | 122.19 | 11 | 11.02 | |
| $S3C_5$ | Ctrl | 100 | 11 | 4.99 | 0.051 |
| | HET | 119.44 | 10 | 12.2 | |
| $S3C_6$ | Ctrl | 100 | 11 | 7.9 | > 0.999 |
| | HET | 104.06 | 11 | 12.85 | |
| $S3C_7$ | Ctrl | 100 | 11 | 7.58 | 0.047 |
| | HET | 145.91 | 11 | 18.23 | |
| S3E | CAG-GFP+Flag-20μm | 8.79 | 14 | 0.76 | Two-Way ANOVA, F (2, 37) = 1.621, P = 0.211 |
| | CAG-GFP+TAK1-20μm | 11.33 | 12 | 0.79 | |
| | CAG-GFP+TAK1$^{KD}$-20μm | 8.93 | 14 | 0.9 | |
| | CAG-GFP+Flag-40μm | 9.86 | 14 | 1.23 | |
| | CAG-GFP+TAK1-40μm | 13.75 | 12 | 1.48 | |
| | CAG-GFP+TAK1$^{KD}$-40μm | 9.36 | 14 | 1.2 | |

| | | | | |
|---|---|---|---|---|
| | CAG-GFP+Flag-60μm | 9.79 | 14 | 1.33 |
| | CAG-GFP+TAK1-60μm | 13.08 | 12 | 2.7 |
| | CAG-GFP+Flag-TAK1$^{KD}$ -60μm | 9.29 | 14 | 1.46 |
| | CAG-GFP+Flag-80μm | 8.36 | 14 | 1.39 |
| | CAG-GFP+TAK1-80μm | 11.83 | 12 | 2.89 |
| | CAG-GFP+TAK1$^{KD}$-80μm | 8 | 14 | 1.46 |
| | CAG-GFP+Flag-100μm | 8.62 | 13 | 1.38 |
| | CAG-GFP+TAK1--100μm | 8.33 | 12 | 2.07 |
| | CAG-GFP+TAK1$^{KD}$-100μm | 7.43 | 14 | 1.22 |
| | CAG-GFP+Flag-120μm | 8.62 | 13 | 1.02 |
| | CAG-GFP+TAK1 -120μm | 8.83 | 12 | 2.14 |
| | CAG-GFP+TAK1$^{KD}$-120μm | 6.46 | 13 | 0.97 |
| | CAG-GFP+Flag-140μm | 9.15 | 13 | 1.22 |
| | CAG-GFP+TAK1-140μm | 10.08 | 12 | 2.96 |

| | | | | |
|---|---|---|---|---|
| | CAG-GFP+TAK1$^{KD}$-140μm | 5.54 | 13 | 1.02 |
| | CAG-GFP+Flag-160μm | 8.15 | 13 | 1.04 |
| | CAG-GFP+TAK1-160μm | 9.58 | 12 | 2.55 |
| | CAG-GFP+TAK1$^{KD}$-160μm | 4.62 | 13 | 0.98 |
| | CAG-GFP+Flag-180μm | 7.31 | 13 | 1.35 |
| | CAG-GFP+TAK1-180μm | 7.75 | 12 | 1.67 |
| | CAG-GFP+TAK1$^{KD}$-180μm | 4.62 | 13 | 0.9 |
| | CAG-GFP+Flag-200μm | 5.82 | 11 | 1.11 |
| | CAG-GFP+TAK1-200μm | 7.3 | 10 | 2.03 |
| | CAG-GFP+TAK1$^{KD}$-200μm | 4.58 | 12 | 1.12 |
| | CAG-GFP+Flag-220μm | 5.36 | 11 | 1.11 |
| | CAG-GFP+TAK1-220μm | 7.75 | 8 | 2 |
| | CAG-GFP+TAK1$^{KD}$-220μm | 3.55 | 11 | 0.85 |
| | CAG-GFP+Flag-240μm | 3.1 | 10 | 0.87 |

| | | | | | |
|---|---|---|---|---|---|
| | CAG-GFP+TAK1-240μm | 8.43 | 7 | 2.15 | |
| | CAG-GFP+TAK1$^{KD}$-240μm | 3 | 9 | 0.58 | |
| | CAG-GFP+Flag-260μm | 3.67 | 6 | 0.99 | |
| | CAG-GFP+TAK1-260μm | 5.86 | 7 | 1.88 | |
| | CAG-GFP+TAK1$^{KD}$-260μm | 3 | 4 | 0.82 | |
| | CAG-GFP+Flag-280μm | 4.8 | 5 | 0.97 | |
| | CAG-GFP+TAK1-280μm | 4.29 | 7 | 1.44 | |
| | CAG-GFP+TAK1$^{KD}$-280μm | 4 | 3 | 1.53 | |
| S3F$_2$ | DMSO | 8.87 | 24 | 0.46 | 0.008 |
| | 5Z-7 | 7.25 | 15 | 0.48 | |
| S3G$_2$ | DMSO-20μm | 10.11 | 9 | 0.75 | 0.561 |
| | 5Z-7-20μm | 10.2 | 5 | 1.11 | |
| | DMSO-40μm | 13.33 | 9 | 1.01 | 0.636 |
| | 5Z-7-40μm | 15.6 | 5 | 2.77 | |
| | DMSO-60μm | 14.67 | 9 | 2.57 | 0.497 |
| | 5Z-7-60μm | 16.4 | 5 | 2.94 | |
| | DMSO-80μm | 13.56 | 9 | 2.2 | 0.975 |
| | 5Z-7-80μm | 13.4 | 5 | 2.09 | |
| | DMSO-100μm | 9.67 | 9 | 1.34 | 0.626 |
| | 5Z-7-100μm | 11 | 5 | 1.95 | |
| | DMSO-120μm | 11.56 | 9 | 1.12 | 0.677 |
| | 5Z-7-120μm | 10.4 | 5 | 2.23 | |
| | DMSO-140μm | 9.67 | 9 | 1.49 | 0.927 |
| | 5Z-7-140μm | 9.2 | 5 | 1.56 | |

| | | | | | |
|---|---|---|---|---|---|
| | DMSO-160μm | 10.33 | 9 | 1.82 | 0.412 |
| | 5Z-7-160μm | 12.4 | 5 | 2.01 | |
| | DMSO-180μm | 12.88 | 8 | 2.61 | 0.699 |
| | 5Z-7-180μm | 14 | 5 | 2.32 | |
| | DMSO-200μm | 10.88 | 8 | 1.68 | 0.911 |
| | 5Z-7-200μm | 12.8 | 5 | 3.61 | |
| | DMSO-220μm | 12.38 | 8 | 2.11 | 0.331 |
| | 5Z-7-220μm | 9.6 | 5 | 2.73 | |
| | DMSO-240μm | 11 | 8 | 1.71 | 0.143 |
| | 5Z-7-240μm | 6.5 | 4 | 1.5 | |
| | DMSO-260μm | 12.5 | 8 | 2.69 | 0.776 |
| | 5Z-7-260μm | 11.33 | 3 | 5.46 | |
| | DMSO-280μm | 9.63 | 8 | 1.19 | 0.988 |
| | 5Z-7-280μm | 10.33 | 3 | 2.96 | |
| $S3H_1$ | TAK1 | 1013.03 | 877 | 55.4 | < 0.001 |
| | $TAK1^{KD}$ | 2103.14 | 854 | 94.27 | |
| $S3H_2$ | TAK1 | 31.35 | 877 | 0.75 | < 0.001 |
| | $TAK1^{KD}$ | 17.63 | 854 | 0.32 | |
| S4F | $GluN1^{WT}$ | 6.38 | 11 | 0.88 | ANOVA P = 0.603, F (2, 33) = 0.5129, P_ $GluN1^{WT}$ vs. $GluN1^{S889A}$ = 0.670, P_ $GluN1^{WT}$ vs. $GluN1^{S889D}$ = 0.642, P_$GluN1^{S889A}$ vs. $GluN1^{S889D}$ = 0.998 |
| | $GluN1^{S889A}$ | 7.29 | 9 | 0.57 | |
| | $GluN1^{S889D}$ | 7.22 | 16 | 0.57 | |
| S4G | $PSD95^{WT}$ | 5.65 | 12 | 0.31 | ANOVA P = 0.578, F (2, 39) = 0.556, P_ $PSD95^{WT}$ vs.$PSD95^{S585A}$ = 0.739, P_ $PSD95^{WT}$ vs. $PSD95^{S585D}$ = 0.560, P_$PSD95^{S585A}$ vs. $PSD95^{S585D}$ = 0.958 |
| | $PSD95^{S585A}$ | 6.27 | 14 | 0.71 | |
| | $PSD95^{S585D}$ | 6.48 | 16 | 0.53 | |
| S4H | $CDKL5^{WT}$ | 7.8 | 10 | 0.82 | ANOVA P = 0.548, F (2, 26) = 0.616, P_ $CDKL5^{WT}$ vs. $CDKL5^{T375A/S377A}$ = 0.522, P_ $CDKL5^{WT}$ vs. $CDKL5^{T375D/S377D}$ = 0.790, |
| | $CDKL5^{T375A/S377A}$ | 6.61 | 9 | 0.8 | |
| | $CDKL5^{T375D/S377D}$ | 7.11 | 10 | 0.64 | |

| | | | | | |
|---|---|---|---|---|---|
| | | | | | P_CDKL5$^{T375A/S377A}$ vs. CDKL5$^{T375D/S377D}$ = 0.890 |
| S4I | CACNG3$^{WT}$ | 7.53 | 4 | 0.32 | ANOVA P = 0.208, F (2, 17) = 1.727, P_ CACNG3$^{WT}$ vs. CACNG3$^{S302A}$ = 0.535, P_ CACNG3$^{WT}$ vs. CACNG3$^{S302D}$ = 0.913, P_CACNG3$^{S302A}$ vs. CACNG3$^{S302D}$ = 0.188 |
| | CACNG3$^{S302A}$ | 6.18 | 7 | 0.94 | |
| | CACNG3$^{S302D}$ | 8.01 | 9 | 0.63 | |
| S4J$_1$ | GluN2B$^{WT}$ | 337.13 | 3917 | 10.2 | K-S test <0.001 |
| | GluN2B$^{S920A}$ | 664.37 | 2739 | 16.13 | |
| S4J$_2$ | GluN2B$^{WT}$ | 14.12 | 3917 | 0.16 | K-S test <0.001 |
| | GluN2B$^{S920A}$ | 16.63 | 2739 | 0.28 | |
| S4K$_1$ | GluN2B$^{WT}$+5Z-7 | 1284.14 | 1126 | 63.33 | K-S test <0.001 |
| | GluN2B$^{S920D}$+5Z-7 | 351.27 | 2074 | 13 | |
| S4K$_2$ | GluN2B$^{WT}$+5Z-7 | 16.85 | 1126 | 0.21 | K-S test <0.001 |
| | GluN2B$^{S920D}$+5Z-7 | 21.79 | 2074 | 0.31 | |
| S5A | ShNC (+) | 1.38 | 8 | 0.23 | ANOVA P = 0.924, F (3, 28) = 0.157 P_ shNC-mCherry vs. shNC-no mCherry = 0.938, P_ shNC-mCherry vs. shTAK1- mCherry = 0.954, P_shNC-mCherry vs. shTAK1-no mCherry = 0.938, P_ shNC-no mCherry vs. shTAK1-mCherry > 0.999, P_ shNC-no mCherry vs. shTAK1-no mCherry > 0.999, P_ shTAK1-mCherry vs. shTAK1-no mCherry > 0.999 |
| | ShNC (-) | 1 | 8 | 0.21 | |
| | ShTAK1 (+) | 1.04 | 8 | 0.54 | |
| | ShTAK1 (-) | 1 | 7 | 0.69 | |
| S5B | tdTomato (+) | 0.77 | 8 | 0.16 | ANOVA P=0.701, F (5, 39) = 0.599, P_ Ctrl-dTomato vs. Ctrl-No-dTomato = 0.961, P_ |
| | tdTomato (-) | 1 | 8 | 0.21 | |
| | tdTomato-TAK1 (+) | 1.11 | 8 | 0.16 | |

| | | | | | |
|---|---|---|---|---|---|
| | tdTomato-TAK1 (-) | 1 | 8 | 0.1 | TAK1-dTomato vs. TAK1-No-dTomato = 0.996, P_ $TAK1^{KD}$-dTomato vs. TAK1KD-No-dTomato = 0.949 |
| | tdTomato-$TAK1^{KD}$ (+) | 0.72 | 8 | 0.27 | |
| | tdTomato-$TAK1^{KD}$ (-) | 1 | 5 | 0.3 | |
| S5C | DMSO | 18.67 | 43 | 0.96 | 0.183 |
| | Nifedipine | 16.56 | 34 | 1.13 | |
| S5D | DMSO | 20.88 | 59 | 1.14 | Wilcoxon matched-pairs signed rank test P < 0.001 |
| | Ifen (3 μM) | 15.68 | 59 | 0.97 | |
| S5E | DMSO | 13.94 | 53 | 1.17 | Wilcoxon matched-pairs signed rank test P = 0.003 |
| | RO (0.5 μM) | 9.57 | 53 | 0.78 | |
| S6A | GFP | 1 | 3 | 0.3 | 0.519 |
| | TAK1-GFP | 1.26 | 3 | 0.21 | |
| S6B | GFP | 1 | 3 | 0.17 | 0.1 |
| | TAK1-GFP | 3.54 | 3 | 0.85 | |
| S6C | GFP | 20.69 | 8 | 1.67 | 0.721 |
| | TAK1-GFP | 20.52 | 8 | 1.78 | |
| $S6E_2$ | GFP-25ms | 1.53 | 28 | 0.03 | unpaired t-test with Welch correction, P_GFP-25 ms vs. TAK1-GFP-25 ms = 0.647, P_GFP-50 ms vs. TAK1-GFP-50 ms = 0.87, P_GFP-100 ms vs. TAK1-GFP-100 ms = 0.297, P_GFP-250 ms vs. TAK1-GFP-250 ms = 0.568, P_GFP-500 ms vs. TAK1-GFP-500 ms = 0.422 |
| | TAK1-GFP-25ms | 1.5 | 32 | 0.04 | |
| | GFP-50ms | 1.6 | 28 | 0.03 | |
| | TAK1-GFP-50ms | 1.59 | 32 | 0.03 | |
| | GFP-100ms | 1.48 | 28 | 0.02 | |
| | TAK1-GFP-100ms | 1.45 | 32 | 0.03 | |
| | GFP-250ms | 1.18 | 28 | 0.02 | |
| | TAK1-GFP-250ms | 1.16 | 32 | 0.02 | |
| | GFP-500ms | 1.05 | 28 | 0.01 | |
| | TAK1-GFP-500ms | 1.07 | 32 | 0.01 | |
| $S6F_2$ | GFP-0.025 mV | 93.82 | 25 | 14.66 | unpaired t-test with Welch correction, P_GFP-0.025 mV vs. TAK1-GFP-0.025 mV = 0.252, P_GFP-0.05 mV vs. TAK1-GFP-0.05 mV = 0.09, P_GFP-0.075 mV vs. TAK1-GFP-0.075 mV = 0.149, P_GFP-0.1 |
| | TAK1-GFP-0.025 mV | 73.45 | 32 | 9.62 | |
| | GFP-0.05 mV | 620.03 | 25 | 55.4 | |
| | TAK1-GFP-0.05 mV | 500.23 | 32 | 41.71 | |

| | | | | | |
|---|---|---|---|---|---|
| | GFP-0.075 mV | 798.37 | 25 | 63.17 | mV vs. TAK1-GFP-0.1 mV = 0.782, P_GFP-0.125 mV vs. TAK1-GFP-0.125 mV = 0.697, P_GFP-0.15 mV vs. TAK1-GFP-0.15 mV = 0.81, P_GFP-0.175 mV vs. TAK1-GFP-0.175 mV = 0.84 |
| | TAK1-GFP-0.075 mV | 678.96 | 32 | 51.57 | |
| | GFP-0.1 mV | 831.95 | 20 | 53.72 | |
| | TAK1-GFP-0.1 mV | 808.57 | 27 | 64.32 | |
| | GFP-0.125 mV | 888.86 | 13 | 71.07 | |
| | TAK1-GFP-0.125 mV | 845.45 | 18 | 84.22 | |
| | GFP-0.15 mV | 867.74 | 8 | 92.31 | |
| | TAK1-GFP-0.15 mV | 834.86 | 12 | 97.91 | |
| | GFP-0.175 mV | 982.45 | 10 | 95.73 | |
| | TAK1-GFP-0.175 mV | 883.63 | 13 | 77.33 | |

All statistical tests were Two-sided Mann–Whitney U test was used unless otherwise stated.
